# TrioNsight: Building a meta-predictor to evaluate the clinical impact of TrioN-like Dbl-homology domain variants

**DOI:** 10.64898/2026.07.17.739111

**Authors:** Alperen Taciroğlu, Yeşim Aydın Son, Andrew C.R. Martin, Christine A. Orengo

**Affiliations:** Department of Health Informatics, Graduate School of Informatics, METU, Ankara, 06800, Türkiye; University College London, London, United Kingdom; Neuroscience and Neurotechnology Center of Excellence, ODTÜ-NÖROM, Ankara, 06560, Türkiye

## Abstract

TRIO is a member of the Rho family of guanine nucleotide exchange factors (Rho GEFs), which promote the exchange of GDP for GTP to activate Rho GTPases and serve as key regulators of cellular signalling pathways. Mutations in TRIO are associated with neurodevelopmental disorders, including intellectual disability and autism spectrum disorders.

TRIO contains two GEF units: one N-terminal and one C-terminal, each of which contains a Dbl-homology (DH) domain that drives its GEF activity to activate Rho GTPases. While the human proteome contains 70 highly conserved DH domains, the N-terminal DH domain of TRIO (TrioN) contains one-third of all reported DH domain pathogenic variants and has many variants of unknown significance.

Numerous variant impact prediction tools exist, but most lack gene-specific considerations. Here, we describe TrioNsight, a meta-predictor designed to predict mutation impacts for TrioN and 12 highly similar human DH domains, including TrioC. TrioNsight exploits the naïve-Bayes algorithm and leverages structural, evolutionary, and physiochemical features of approximately 1500 highly similar DH domains (TrioN-like DH domains) from 294 species. TrioNsight surpasses all available predictors, including AlphaMissense, achieving a Matthews’ Correlation Coefficient of 0.890. Additionally, we provide a variant impact map that details the impacts of mutations at each position in the DH domain of these proteins, which can be valuable for clinical assessments. Furthermore, our approach establishes a standardised workflow adaptable for creating domain-specific variant predictors for other protein families, offering a template for improved variant interpretation.

## Introduction

The TRIO family of proteins is evolutionarily highly conserved, with some well-characterised members including the vertebrate paralogs TRIO and KALRN, and the invertebrate orthologs *Drosophila* TRIO protein, and *C.elegans* UNC-73 protein [1,2]. These proteins are involved in multiple biological processes, such as developing nervous systems and actin remodelling coordination necessary for cell migration and growth [1,3,4]. Interactions with diverse biological molecules such as cytoskeleton-interacting proteins, kinases, and Rho family GTPases (Rho GTPases) enable these diverse roles [5].

The ‘*triple-functional domain protein*’ (TRIO) contains two guanine nucleotide exchange factor (GEF) units and a serine/threonine kinase domain at the C-terminal end. All four characterised TRIO family proteins contain two Dbl-homology (DH) domains, with one in each GEF unit. These domains represent a specialised subtype within the RhoGEF domain family that regulate Rho GTPases by catalysing the exchange of GDP for GTP [6]. TRIO and KALRN are the only proteins in the human proteome with two DH domains [4], named the ‘N-terminal DH domain’ and the ‘C-terminal DH domain’ [3]. TRIO is highly conserved throughout evolution and therefore highly intolerant to loss-of-function variants [7–9]. Figure 1 presents the multi-domain architecture of TRIO and the structure of its N-terminal DH-PH (plekstrin homology) domain pair.

**Figure 1:**
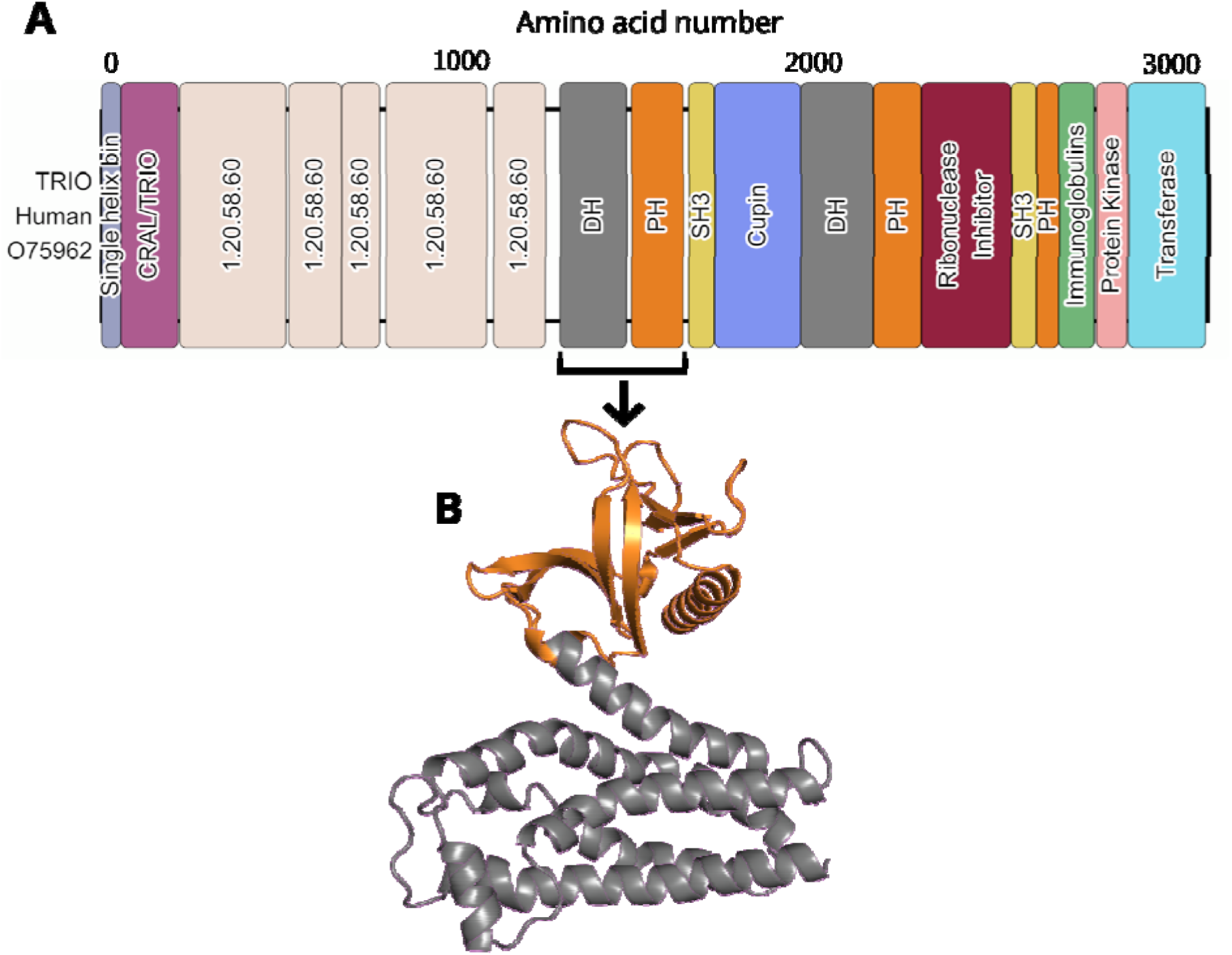
The multi-domain architecture and structure of the N-terminal DH-PH domains of th human TRIO protein (PDB: 1NTY) (RRID:SCR_012820) [23]. A) The multi-domain architecture of the human TRIO protein based on the CATH-Gene3D (RRID:SCR_007672) resource. TRIO contains two tandem DH-PH domains (also called a GEF unit), named as either the N-terminal or C-terminal DH-PH domain based on their location on the protein. B) The structure of the N-terminal DH-PH domain pair of the TRIO protein. The DH domain is coloured in white-grey, and the PH domain is coloured in orange.

Rho GTPases take part in the control of the actin cytoskeleton, leading to processes such as cell migration, cell division, and endocytosis [10]. 81 different RhoGEF domains have been discovered in the human proteome and these can be separated into two subfamilies: the DH domains containing 70 members and the DOCK-family guanine nucleotide exchange factors containing 11 members [11–13]. Here we focus on the more common DH domains which contain a high number of non-synonymous mutations. PH domains almost invariably follow DH domains in the protein sequence [14], although most of the guanine nucleotide exchange activity occurs in the DH domain [15,16]. Variants in DH domains are linked to multiple human diseases including Aarskog–Scott syndrome (AAS) [17], primary immunodeficiency diseases (PIDs) [18], cancers [19], and neurodevelopmental disorders [7,20–22]. To understand the molecular function of DH domains and the clinical impact of DH domain variants better, we developed a specialised variant impact predictor tailored specifically for the N-terminal DH domain of TRIO (TrioN) and other TrioN-like DH domains, including the C-terminal DH domain of TRIO (TrioC).

The TRIO N-terminal DH domain (TrioN) is densely populated with nonsynonymou variants (see Supplementary Figure S1). A small percentage of these variants have been classified as pathogenic or neutral in ClinVar (RRID:SCR_006169) [24], GnomAD (RRID:SCR_014964) [25], and HGMD [26] based on clinical verification and minor allele frequency. The majority, however, fall under the category of ‘unknown significance’. Numerou variant impact prediction (VIP) algorithms have been developed, and we assessed the performance of automated VIP algorithms on the variants located in TrioN. Previous reports state that automated variant impact prediction algorithms, trained on millions of variants across the human proteome, have varying performance on individual protein families [27,28]. Considering their functional and clinical importance, we decided to develop a variant impact prediction algorithm, TrioNsight, specifically for DH domains. TrioNsight is trained and tested on approximately 1500 highly similar DH domains (‘TrioN-like DH domains’) from 294 species.

Pathogenicity predictions of four established predictors were used as component predictors in training TrioNsight together with additional sequence and structural analyses. Thus, TrioNsight is an ‘enhanced meta-predictor’ (a model that uses predictions from other predictors together with additional features). The component predictors are all recently-developed methods that have been benchmarked by comparing with the best-performing methods in the latest critical assessment of genome interpretation (CAGI-6) [29], namely: VariPred [30], AlphaMissense [31], PHACT [32] and PHACTboost [33]. Recently-developed methods were preferred because of the fast-paced developments in the variant impact prediction field. A summary of these component predictors is provided in the Supplementary Section ‘<u>Component predictors of</u> <u>TrioNsight</u>’.

TrioNsight was benchmarked against all of its component predictors, and other highly cited and commonly used predictors such as ESM-1b [34], CADD (RRID:SCR_018393) [35], MutPred2 [36], and PolyPhen-2 (RRID:SCR_013189) [37].

TrioNsight outperforms its closest competitor in Precision, Accuracy, False Positive Rate, F1, and MCC metrics, while demonstrating competitive performance in Recall and False Negative Rate metrics. Beyond these robust results for TRIO and related DH domains, our approach establishes a standardised framework for developing domain-specific variant impact predictors. This methodology provides a template that can be systematically applied to other protein domain families where variant interpretation remains challenging.

## Materials and methods

### Selection of TrioN-like DH domains

TrioNsight learns the likelihood of variants in targeted DH domains being pathogenic from a set of sequences (named the ‘TrioN-like DH dataset’). This dataset was compiled using a stringent protocol to select appropriate DH domains from those identified in CATH-Gene3D [38]. The process comprised a sequence-based analysis using functional families generated by the FunFams algorithm [39], and a structure-based analysis using experimental and AlphaFold2- generated (RRID:SCR_025454) [40] models. It aimed to assemble a diverse set of domain sequences from as many organisms as possible, while ensuring that the selected sequences have a high probability of being structurally and functionally similar to TrioN (see Figure 2), and thereby expanding the dataset of pathogenic and benign variants required to train the TrioNsight predictor.

**Figure 2:**
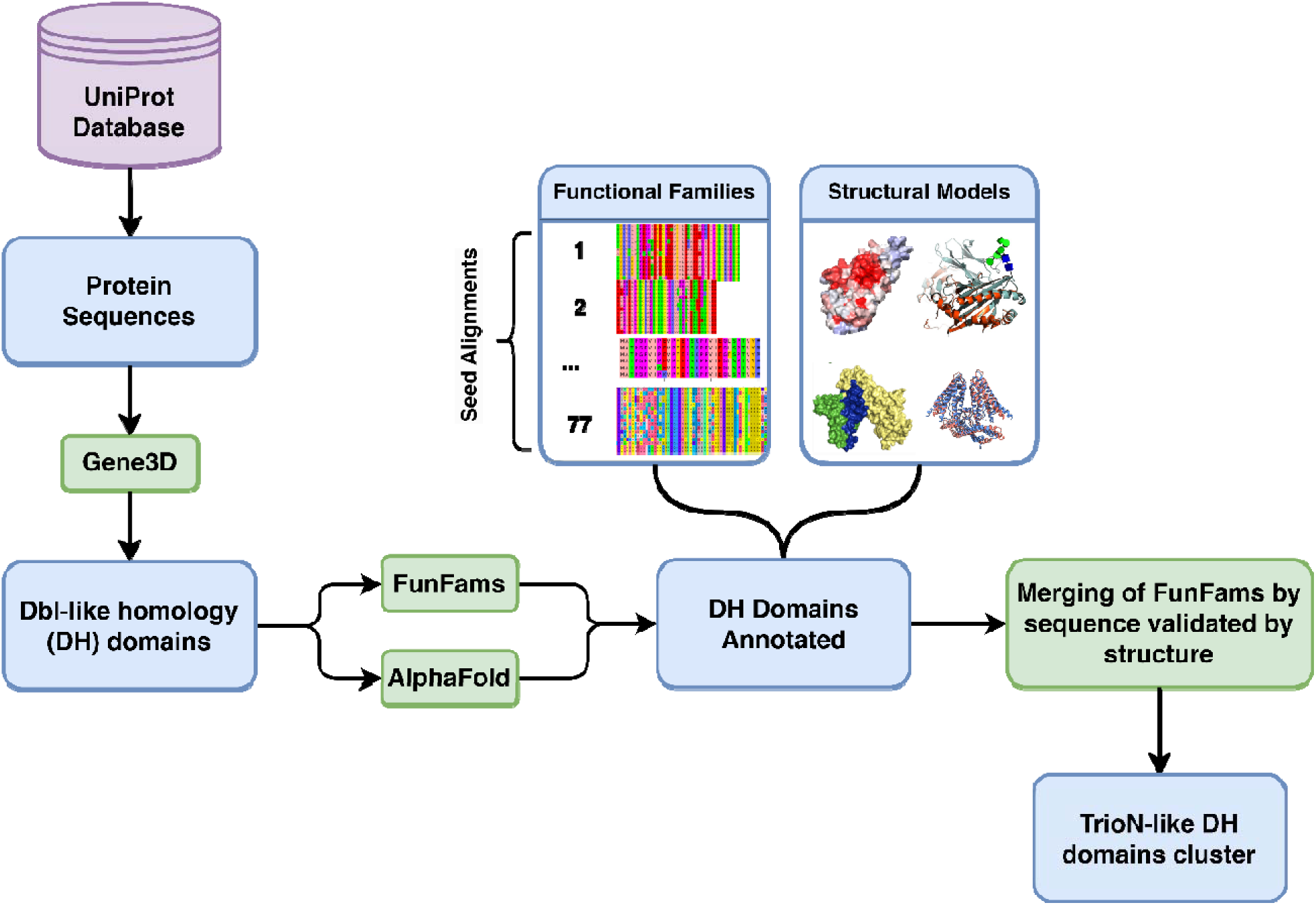
Workflow to obtain TrioN-like Dbl-homology (DH) domains for TrioNsight. DH domains were identified using CATH-Gene3D [38] domain annotations of the protein sequences from UniProtKB (RRID:SCR_004426) [41]. Structural models were created for each of these DH domains using AlphaFold2. DH domains were assigned to their corresponding functional families using the FunFams algorithm [39]. Subsequently, HHalign (RRID:SCR_016133) [42] was used to identify highly similar FunFams using an e-value threshold of 10^−24^ to generate a larger set of sequences containing the TrioN DH domain. Pairs of FunFams were merged if the relatives were structurally similar. This was determined using structure-based analyses, namely the CATH-SSAP [43] structure comparison method with thresholds of 80% residue overlap and RMSD < 2.5 Å superposition against the reference structure of the TrioN DH domain (see text). This gave a broad set of sequences that contained the TrioN DH domain, but in which relatives were structurally highly similar and likely to be functionally similar. The final subcluster containing TrioN was named the ‘TrioN-like DH domains cluster’. It contains DH domains from FF-000001, FF-000002, FF-000008, FF-000019, and FF-000028 functional families of the CATH superfamily 1.20.900.10.

Full details of the selection process are provided in the Supplementary Section ‘Extended methodology for the selection of TrioN-like DH domains’.

### Conservation analysis and variant mapping of TrioN-like DH domains

We identified conserved functional residues by performing conservation analysis on the DH superfamily, the specific FunFam containing TrioN, and the curated TrioN-like dataset. We then mapped nonsynonymous variants from ClinVar, GnomAD, and HGMD Pro to these domains, after applying filtering criteria to categorise them as pathogenic or benign. Full details of the alignment protocols, conservation scoring, and variant curation are provided in the Supplementary Section ‘<u>Extended methodology for the conservation analysis and variant</u> <u>mapping of TrioN-like DH domains</u>’.

### Inheriting variants from one TrioN-like DH domain to another

Increasing the data size used for training is crucial both to achieve higher performance and to prevent overfitting [44,45]. ‘Variant inheritance’ (also called ‘variant aggregation’) was used to increase the amount of training data with which to train TrioNsight. The aggregation of variants is based on earlier work that detects mutational enrichment [46–48].

AlphaFold2 models (see Supplementary Section ‘Creating AlphaFold models for the TrioN-like DH domains’), and experimental structures from the PDB, were aligned using a local installation of mTM-align [49], which generates a multiple structure alignment progressively based on pairwise structure alignments.

Variants from other human proteins within the TrioN-like DH dataset were projected onto the equivalent structural position of each human DH domain in the dataset, using the multiple structure alignment positions as a guide (see Figure 3). This was only carried out if the native residues in the two structures being aligned were identical at that position. Since the TrioN-like DH dataset domains share similar structures and functions, inherited variants are likely to be involved in similar functional roles.

**Figure 3:**
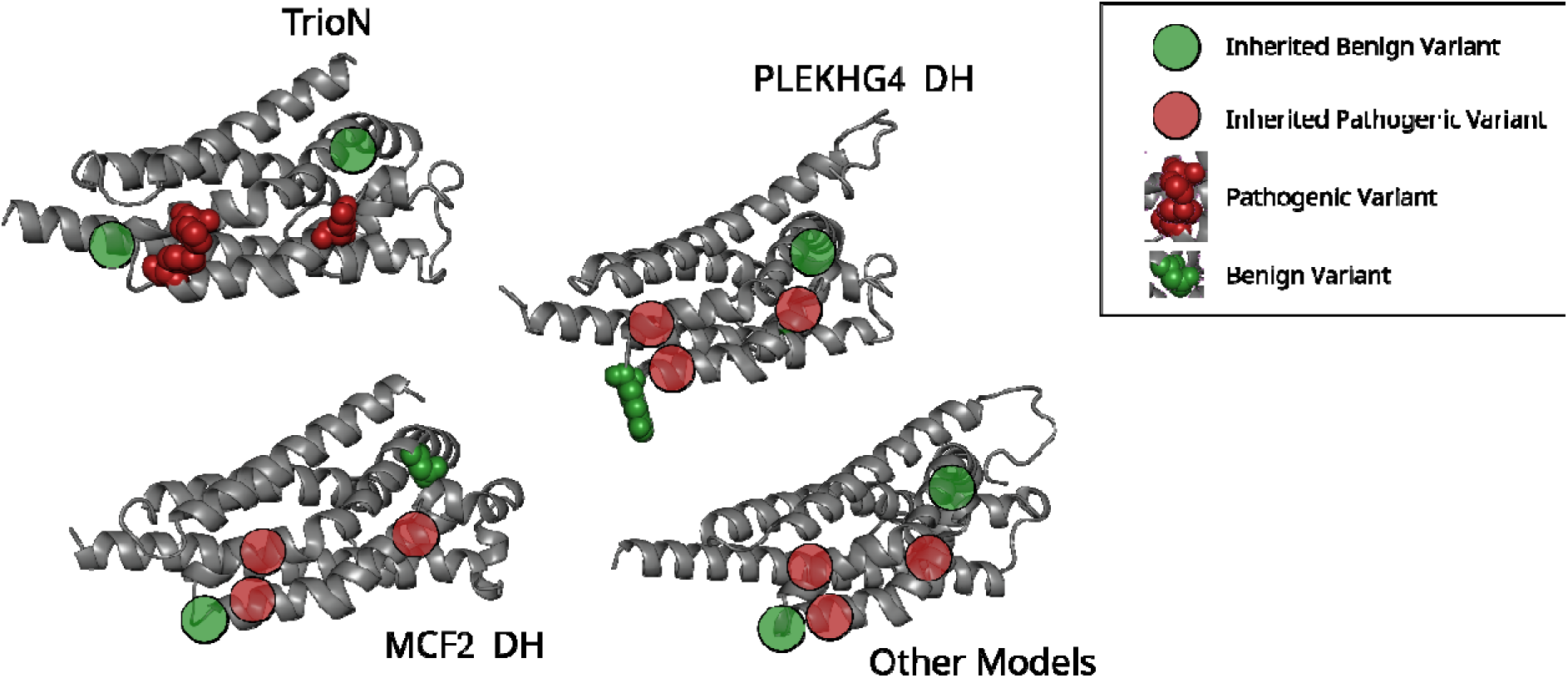
Inheriting variants in TrioN-like DH domains using the structural alignment as a guide to enable mutational enrichment. The procedure effectively increased the number of data points with which to train the model. Variants were only inherited if, in the multiple structure alignment, the native residue in the donor sequence is identical to the native residue in the inheriting sequence.

### Variant clustering and deriving Distance-to-Cluster features

We calculated the Distance to Variant Clusters (DVC) features based on previous work that applied 3D-clustering of variants where there had been some evidence of positions of mutations being linked to different pathogenic phenotypes [50]. To calculate the DVC, all variants located on the TrioN-like DH domains were mapped to a representative TRIO structure (PDB: 1NTY [23]) using the multiple structural alignment described above. 3D clustering of variants was then performed using strucclus [50] which simply clusters C-alpha positions of mutated residues using single linkage hierarchical clustering. A range of numbers of clusters are assessed using a chi-squared test to identify the number of clusters that shows the most significance in separating pathogenic and neutral variants.

### TrioNsight - Features

TrioNsight is an enhanced meta-predictor with 47 unique features which fall into 3 categories (see Figure 4):

- **Sequence-based Analyses:** This category includes BLOSUM80 [51] scores, ScoreCons [52] scores, Grantham distance [53] scores, Functional relevance by Swiss-Prot obtained as a part of the SAAPdap pipeline [54] (2 features), and VarSite [55] conservation scores obtained from the ProtVar tool [56]. (A total of 6 features).
- **Structural Analyses:** This category includes SAAPdap analyses [54] (30 features), distances to variant clusters (5 features), and FoldX (RRID:SCR_008522) [57] (2 features). FoldX results were obtained from the ProtVar tool [56] by supplying the genomic location of the corresponding variants. (A total of 37 features).
- **Variant Prediction Algorithms:** This category incorporates pathogenicity probability predictions from established algorithms: AlphaMissense [31], PHACT [32], PHACTboost [33] and VariPred [30]. (A total of 4 features).

**Figure 4:**
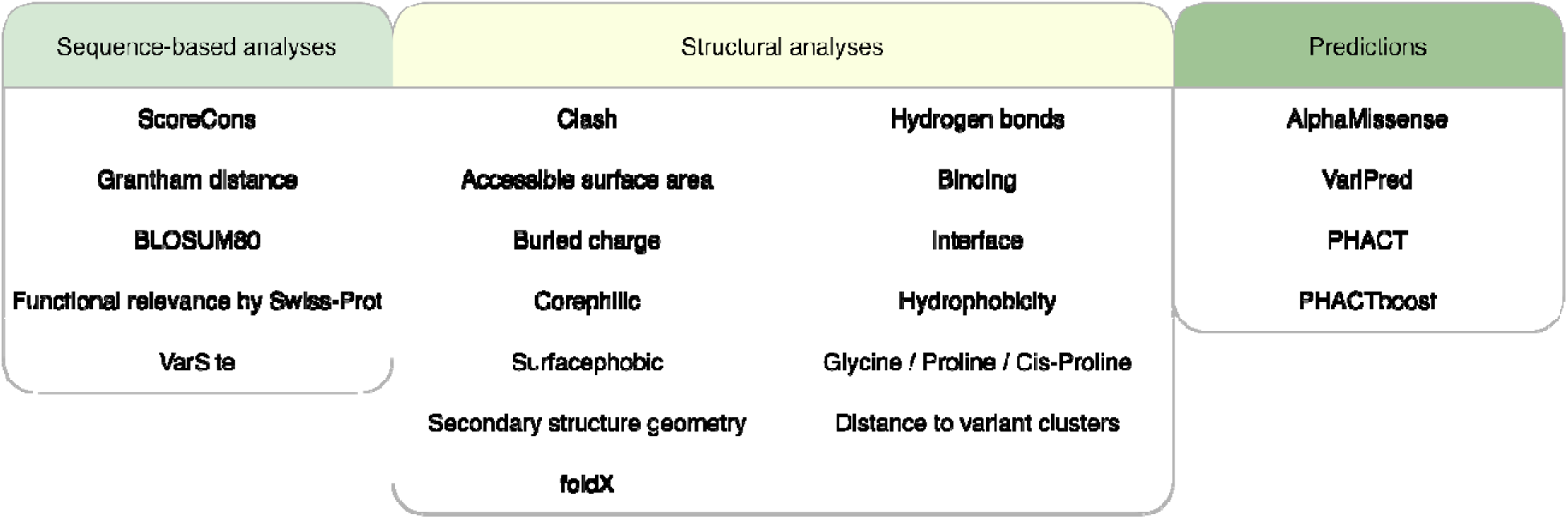
The features and predictions used to train TrioNsight. Scorecons, Grantham distance and BLOSUM80 scoring were obtained by analysing the multiple sequence alignment of the TrioN-like DH domains. SwissProt features are a SAAPdap output. VarSite, FoldX scores were obtained from the ProtVar tool as separate scores for each corresponding variant. Distances to variant clusters were calculated on the five clusters created by the strucclus algorithm on variants both occurring in, and inherited to, TrioN-like DH domains. The remaining structural analyses were performed by running SAAPdap on the structural models. External predictions were obtained by running the variant prediction algorithm on the variant dataset, which returns a probability of pathogenicity and predicted class for AlphaMissense, PHACT and PHACTboost, and the predicted class for VariPred. The predicted class is either pathogenic or benign for VariPred, PHACT and PHACTboost, and pathogenic, benign, or ambiguous for AlphaMissense. The probability of pathogenicity from AlphaMissense, PHACT and PHACTboost algorithms and the predicted class for the VariPred algorithm were used for the training of TrioNsight.

See Supplementary Section ‘Feature generation protocols’ for detailed methods on feature generation.

### Training TrioNsight and optimising the features

TrioNsight was trained using 18 different methods (see Supplementary Table S1) using the ‘train’ function of the caret package [58] in R. Features were centred and scaled during training as part of the ‘train’ function. The leave-one-out train control method was used to ensure that no variant from the same residue was included both in the training set and test set. This was a crucial step of the training because inheriting variants onto multiple structures amplified data points coming from each variant. Mixing these projections in the train and test sets would result in an inflated performance of the predictor.

After the initial training, feature optimisation was implemented through an iterative process. Features that did not contribute to the predictor’s performance were systematically identified and removed, and the models were repeatedly retrained with different feature subsets. For each iteration, performance was evaluated using Matthews’ correlation coefficient (MCC), and this process was continued until maximum MCC values were achieved. The model configuration that yielded the highest performance metrics was selected as the final TrioNsight model (see Figure 5). Full details of the machine learning model selection process are provided in Supplementary Section ‘<u>Model Selection</u>’.

**Figure 5:**
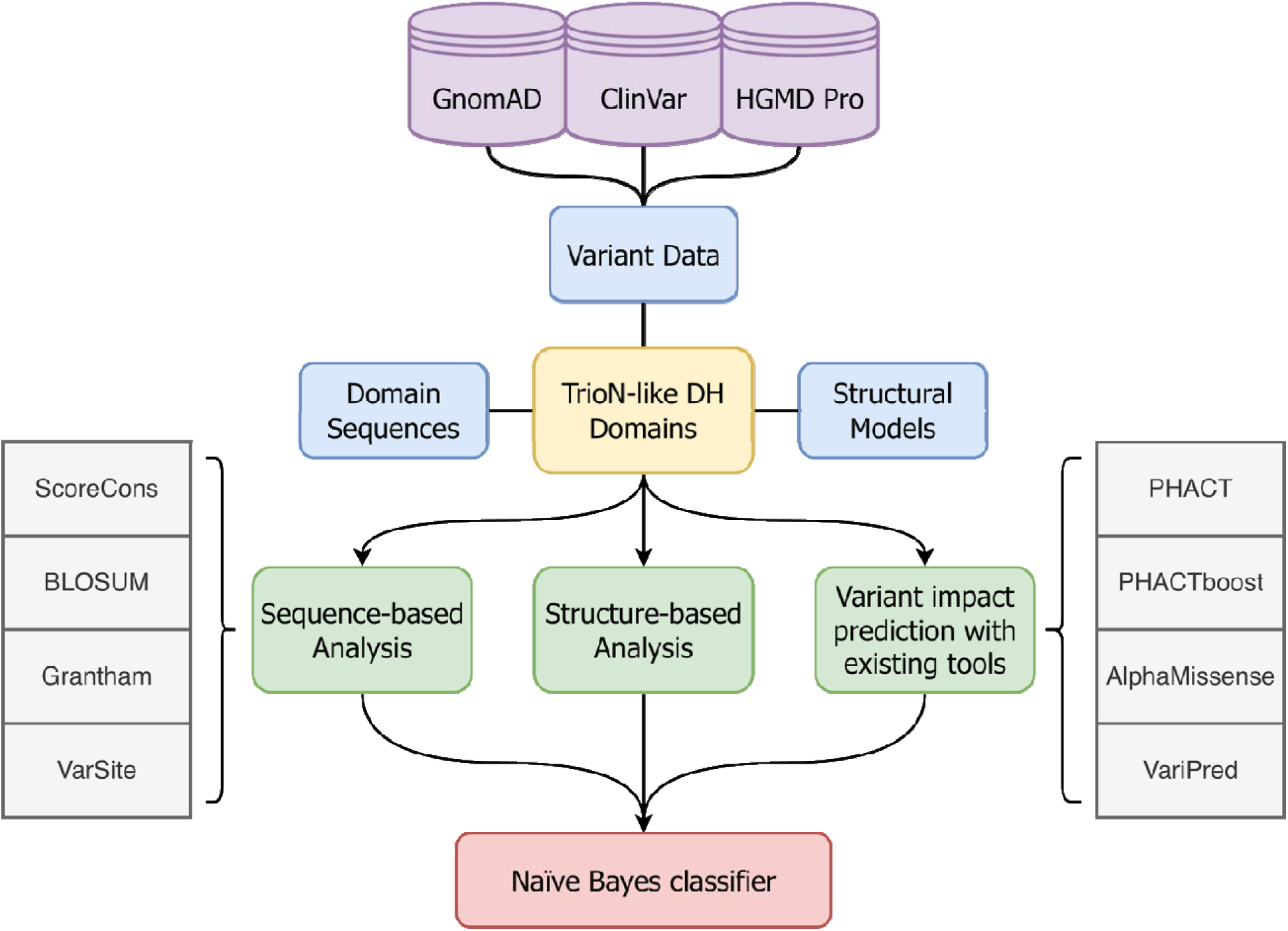
The workflow shows the raw data and the features that were used to train TrioNsight. ScoreCons [52] scoring, BLOSUM [51] scoring, and Grantham distances [53] were obtained from the multiple sequence alignment of the highly similar TrioN-like DH domain sequences. VarSite [55] conservation scores were obtained through the ProtVar API. The SAAPdap algorithm [54] was run on the DH domain models to obtain the structural features. PHACT [32], PHACTboost [33], AlphaMissense [31] and VariPred [30] external prediction algorithms were incorporated into the training of TrioNsight. The optimal set of features was selected by evaluating the performance of the model with various feature subsets and machine learning methods.

### Benchmarking TrioNsight predictions

Pathogenicity predictions from AlphaMissense, VariPred, PHACT, PHACTboost, MutPred2, CADD and PolyPhen-2 were used in benchmarking TrioNsight. Performance was measured for each method using Precision, Recall, Accuracy, False Positive Rate, False Negative Rate, F1 Score and Matthews’ Correlation Coefficient (MCC) metrics. (See Supplementary Section ‘<u>Equations</u>’.)

See Supplementary Section ‘Data circularity tests’ for details of the validation approaches used to prevent data leakage and circular reasoning.

### Variant impact map as TrioNsight model output

A comprehensive variant impact map was created by systematically analysing all possible amino acid substitutions across every residue position in the TrioN-like DH domains. The same feature categories used to train TrioNsight were extracted for each hypothetical variant. Variants were mapped onto multiple structural models representing different homologous proteins in the Trion-like DH dataset. The predicted pathogenicity scores therefore varied depending on the specific structural context. For instance, a single variant might have a pathogenicity probability of 0.60 when evaluated using one structural model but have a pathogenicity probability of 0.40 when assessed using another model. To account for this structural context-dependent variation, the mean pathogenic probability and standard deviation (sd) were calculated for each variant across all applicable structural models. These statistical measures were then used to assign appropriate classification categories as follows:

To classify variant predictions into clinically relevant categories, the following criteria were applied: variants with low prediction variability (σ < 0.05) and high pathogenicity scores ( > 0.75) were classified as ‘Pathogenic’; those with higher variability (σ ≥ 0.05) but high pathogenicity scores as ‘Likely pathogenic’; variants with low variability and low pathogenicity scores ( < 0.25) as ‘Benign’; and those with higher variability but low pathogenicity scores as ‘Likely benign’. All other predictions were classified as ‘Uncertain’, with any prediction showing high variability (σ > 0.2) automatically classified as ‘Uncertain’ regardless of its mean score.

See Supplementary Table S2 for the complete set of 50,122 variant impact predictions with clinical classifications across all residue positions in the TrioN-like DH domains.

## Results and discussion

### Variant localisation analysis justifies the specific focus on DH domains

The N-terminal and C-terminal ends of proteins containing DH domains can harbour regions and domains that are known to interact with the DH domains such as C2, SH2, SH3, PH domains, or interdomain linkers. Of the domains typically associated with human DH domains, only PH domains and SH3 domains were found in the vicinity of the TrioN-like DH domains. We analysed the variant distribution for these neighbouring regions and observed that PH domains were mostly barren in terms of pathogenic nonsynonymous variants (See Supplementary Figure S2). Furthermore, SH3 domains were only sporadically present in the multi-domain architecture of the proteins of interest. Consequently, TrioNsight was trained using DH sequences and variant data from DH domains only.

### Conserved region similarity validates the selection of TrioN-like DH domains and indicates novel functional sites

TrioN was classified in FunFam FF-000001 along with the N-terminal DH domain of KALRN, and DH domains of Proto-oncogene DBL (MCF2) [59], Guanine nucleotide exchange factor DBS (MCF2L) [60], and Probable guanine nucleotide exchange factor MCF2L2 [61]. The C-terminal DH domain of TRIO (TrioC) was classified in FunFam FF-000008 along with KalrnC and ARHGEF25 [62]. To maximise the number of DH domain relatives which can be used to find conserved residues likely to be of functional relevance and to maximise the number of pathogenic and benign variants that can be used in training the TrioNsight predictor, the TrioN-like DH domains dataset was compiled. This final dataset contained domains from multiple functional families. See Supplementary Table S3 for TrioN-like DH domain dataset and Supplementary Figure S3 for multidomain organisation of human proteins that contain a TrioN- like DH domain.

The similarity of these DH domains obtained after the sequence and structural-based protocols was verified by analysing three sets of DH domains i.e. (1) DH all domains, (2) DH FunFam 000001 domains, (3) DH merged FunFam domains (including FF-000008).

The first of these sets contained representatives from all DH domain functional families. The conservation pattern on this broad set of sequences was visualised to serve as a negative control (see Figure 6A). The second set contained highly specific DH domains only from the functional family FF-000001 (see Figure 6B). The enhancement of conserved residues (as measured by Scorecons) can be seen. Subsequently, the conservation patterns for the TrioN-like DH domain dataset were visualised. This is the TrioN-like DH set of DH domains obtained by the protocol using the sequence and structure-based comparisons to merge the FunFams (see Figure 6C) which achieved a high information content (DOPS score of 85.1). See <u>Supplementary</u> <u>Movie S1</u> for rotational visualisation of amino acid conservation across different DH domain sets mapped onto the representative TrioN structure.

**Figure 6:**
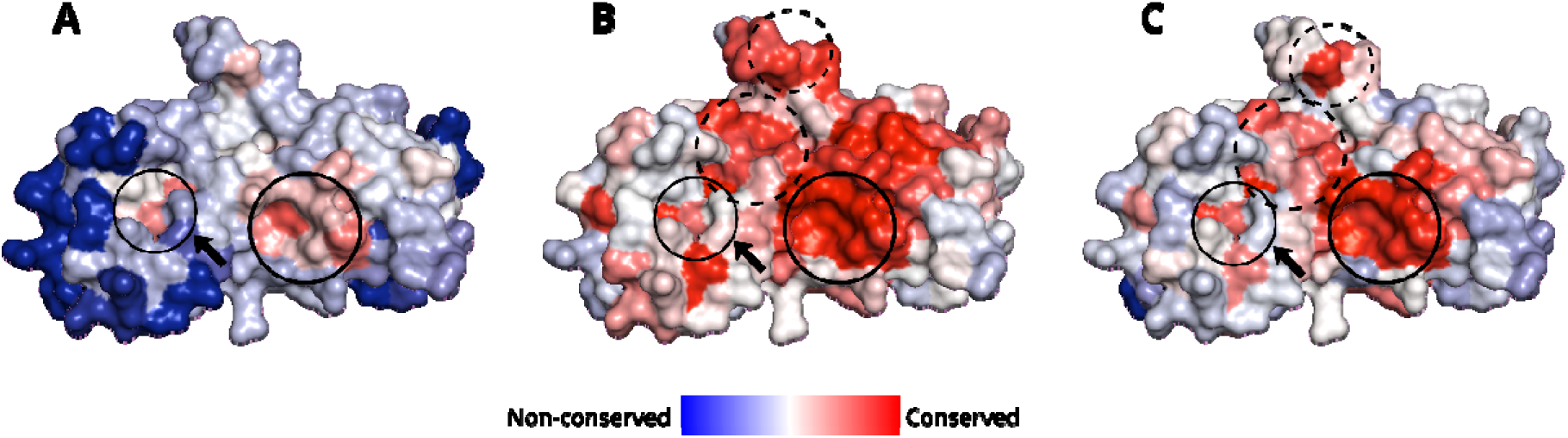
Amino acid conservation of three different sets of DH domains mapped on the representative TrioN structure (1NTY - DH domain coordinates only). Multiple sequence alignment was performed on the three sets of DH domains containing A) non-redundant sequences from all DH domain functional families (selected from clusters of functional family relatives clustered using the MMSeqs2 algorithm with a 70% sequence identity threshold), B) DH domain functional family of TrioN, FF-000001, and C) TrioN-like DH domains obtained by merging FunFams based on sequence and structure similarity. The representative TrioN structure is colour-coded according to Scorecons sequence conservation scores obtained using these multiple sequence alignments. The regions conserved in all three sets are circled using a continuous line. The regions conserved only in two sets are circled using a dashed line. The potential novel functional site is marked with arrows that is conserved across all three sets. The unmarked conserved sites in B are associated with TrioN - Rac1 binding.

Residue conservation patterns within structurally relevant regions were found to be maintained even after the expansion beyond the initial functional family. The FF-000001 functional family DH domains and TrioN-like DH domains share four conserved regions which are likely to be involved in ligand binding. These residues coincide with the TrioN-Rac1 binding interface reported by previous research [23,63]. Furthermore, Rac1 is a known binding partner of GEFs in our list of proteins containing TrioN-like DH domains such as KALRN and PLEKHG2 [64–66].

Of these four conserved regions, two regions are conserved across all three sets. These two regions are likely to be fundamental to the protein’s overall structure and function (see Figure 6). Additionally, a potential novel functional site has been identified within one of the regions conserved across all three sets, corresponding to residues 1340-1344 in TRIO (UniProt ID: O75962), suggesting further functional significance beyond the previously characterised binding interface. The retention of the conserved regions in the set of merged FunFams (TrioN- like DH dataset) implies that the TrioN functional family was expanded robustly with domains likely to be functionally similar, using the dataset selection protocol described in the Methods.

Furthermore, the high structural similarity of human domains within the TrioN-like DH dataset is verified in their superimposed structures as shown in Supplementary Figures S4 + S5, and Supplementary Table S4.

### Variant inheritance increases the amount of training data for predicting variant impacts in homologous domains

Variants from other human proteins within the TrioN-like DH dataset were projected onto each human DH domain in the dataset, guided by the multiple structure alignment. Because the native residue of the domains containing the variant needs to be identical to the native residue in the protein used to inherit the variant, the number of variants that could be inherited in each of the models varied. Despite this prerequisite, variant inheritance expanded 61 data points (from the sum of pathogenic and benign variants) to 1051 data points for structural features, a large increase in the amount of data on which to train the predictor. While any single structural model can only display a subset of these variants (e.g., 29 variants on the TrioN structure in Figure 7) based on residue identity, the full complement of 61 variants is preserved and utilised across the diverse structural contexts of the entire dataset.

**Figure 7:**
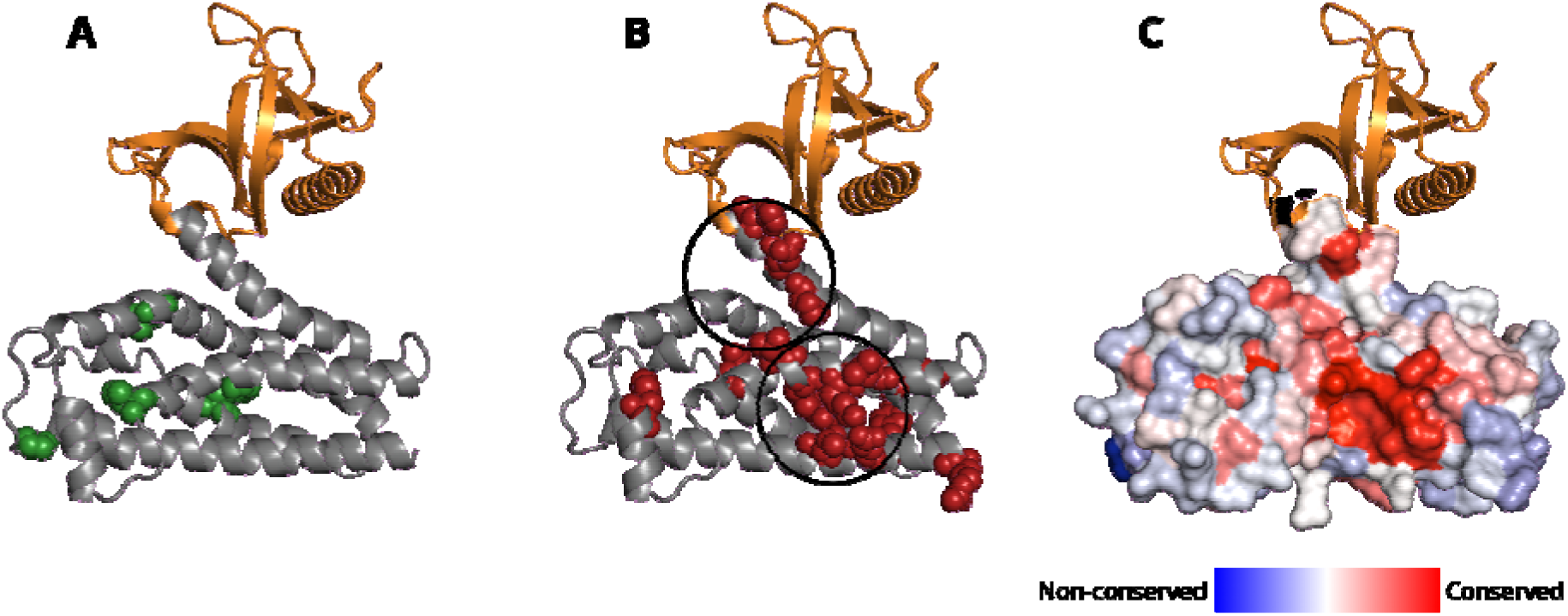
Benign and pathogenic variants located in, or inherited by, the N-terminal DH domain of TRIO visualised side-by-side with the ScoreCons residue conservation scores on th representative experimental TrioN DH-PH model (PDB: 1NTY). The figure shows only the 29 variants out of 61 that could be inherited onto this structure because the native residue is th same, as described in the methods. Variants are colour-coded as green for benign variants and red for pathogenic variants based on their clinical pathogenicity classification. ScoreCons scores are obtained from the multiple sequence alignment of the TrioN-like DH domains. A) Benign variants plotted on the TrioN model (shown as space-filled atoms). B) Pathogenic variants plotted on the TrioN model. C) TrioN model surface view coloured by Scorecons conservation scores.

However, the amount of data coming from sequence-based features, such as Scorecons conservation scores, remains at 61 (i.e. the number of positions mutated), as the information in the multiple sequence alignment used to derive the conservation score does not change.

### Pathogenic variants localise separately from the neutral variants on the TrioN structure

Variants inherited from each of the TrioN-like DH domains and TrioN variants were plotted on the representative TrioN DH-PH model (PDB: 1NTY) together with a surface overlay of residue sites having significant ScoreCons residue conservation (see Figure 7). The 23 pathogenic variants coincide with regions having higher conservation scores overall compared with the 6 neutral variants. Manual inspection revealed regions where pathogenic variant accumulate and neutral variants tend not to occur (see circled regions in Figure 7B).

### Variant clusters on the representative TrioN DH structure

Although DH domains undergo allosteric modulation, our dataset of TrioN-like DH domains have remarkably similar structures owing to the strict selection criteria applied to obtain them. This set is remarkably successful in training the predictor, perhaps suggesting that this conformation is critical for protein function. Functionally important regions are suggested by a high number of pathogenic variants from different proteins clustering in similar spatial locations. Similarly, if benign mutations accumulate in specific areas of the structures, these areas are unlikely to be crucial for the function (see Figure 8). See Supplementary Section ‘<u>Optimising</u> <u>strucclus parameters’</u> for more details.

**Figure 8:**
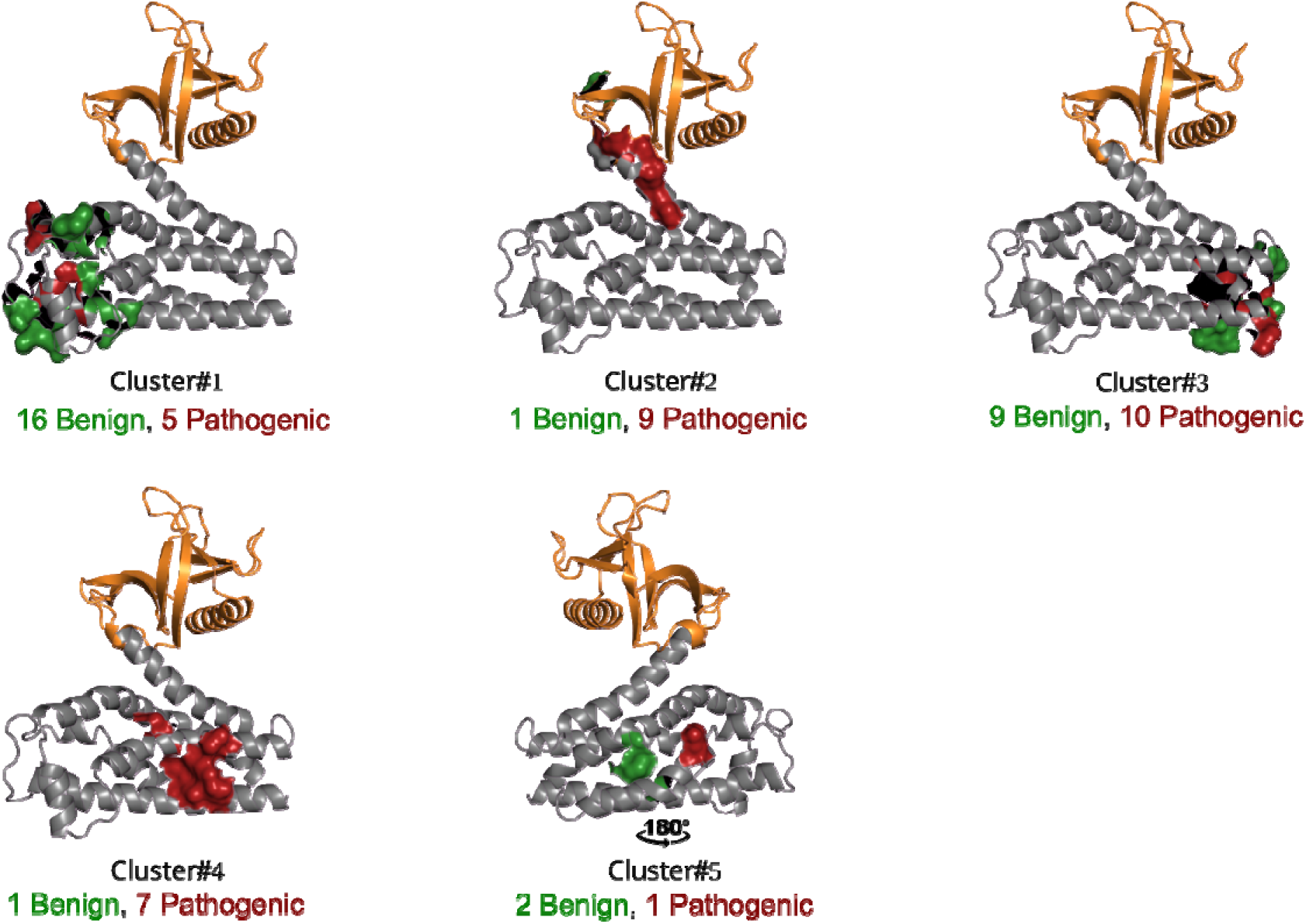
Clustering of the variants on the representative 1NTY TrioN-DH model. Five variant clusters were identified by running the strucclus algorithm on the variants inherited across TrioN-like DH domains. TRIO-Rac1 binding was considered in choosing the optimal number of clusters. The variants forming the clusters are shown space-filled, with red representing pathogenic variants and green representing benign variants.

Cluster#2 and cluster#4 contain the residues involved in Rac1 binding in TRIO. These clusters also have the highest ‘pathogenic variants / all variants’ ratio. (Cluster#2 90% pathogenic; cluster#4 ∼86% pathogenic.) See Supplementary Table S5.

Cluster#1 has the highest number and ratio of benign variants (∼76%). The mutations in the residues covered by this cluster have a lower likelihood of disrupting the overall function and structure of the domain compared with clusters #2 and #4.

Cluster#3 contains a mixture of benign and pathogenic variants. Cluster#5 contains too few variants to draw meaningful conclusions about their importance.

### Comparison of TrioNsight with its closest competitors

The performance of all 18 machine learning methods tested for TrioNsight was measured using Precision, Recall, Accuracy, False Positive Rate, False Negative Rate, F1 Score and MCC metrics. MCC was the preferred measure of performance because it is reliable and intuitive [67], and provides a more comprehensive measure by considering all aspects of the confusion matrix to an equal degree [68,69]. AUC was not used because it integrates performance across the full range of prediction thresholds while MCC uses a fixed threshold as would be required for a real predictor. The naïve-Bayes algorithm was selected as the best-performing method based on these tests and was used for TrioNsight’s predictions.

As described above, pathogenicity probabilities obtained from AlphaMissense, VariPred, PHACT and PHACTboost were used as features in the training of TrioNsight. CADD, ESM-1b, MutPred2 and PolyPhen-2 algorithms were only included as benchmarking methods because they are widely used.

To prevent data circularity, we analysed overlaps between our 61-variant evaluation dataset and the training sets of all component and benchmarking predictors. Overlaps were identified for MutPred2, PHACTboost, and VariPred, whereas CADD, AlphaMissense, and PHACT were largely exempt. To address potential bias, we implemented a filtered benchmarking protocol where overlapping variants were excluded from the evaluation of specific methods. Consequently, TrioNsight was evaluated using a modified leave-one-out cross-validation (LOOCV) approach.

TrioNsight outperforms all benchmark methods by a wide margin based on the MCC metric (0.890) (see Table 1). According to MCC, PHACTboost, MutPred2 and VariPred are TrioNsight’s closest competitors in that order (MCC = 0.775, 0.754 and 0.750 respectively). ESM-1b performs better than TrioNsight in Recall and False Negative Rate metrics, which shows ESM-1b’s ability to predict pathogenic variants correctly, at the expense of identifying benign variants as pathogenic.

**Table 1:** Performance of the predictors used in benchmarking and/or training of TrioNsight were measured after removing the variants found to be overlapping in their respective training dataset and TrioN-like DH dataset. TrioNsight was trained using a modified leave-one-out cross-validation approach where overlapping variants were removed only from the test sets. Performance was assessed across seven metrics: precision, recall, accuracy, false positive rate, false negative rate, F1 score, and Matthews correlation coefficient (MCC). Best-performing scores for each metric are shown in bold. TrioNsight achieved the best performance in five out of seven metrics, including the highest MCC of 0.890, whilst ESM-1b demonstrated superior recall and false positive rate.

| Predictor | Precision | Recall | Accuracy | False Positive Rate | False Negative Rate | F1 Score | MCC |
| --- | --- | --- | --- | --- | --- | --- | --- |
| CADD | 0.871 | 0.844 | 0.853 | 0.138 | 0.156 | 0.857 | 0.705 |
| ESM-1b | 0.775 | **0.969** | 0.836 | 0.310 | **0.031** | 0.861 | 0.692 |
| AlphaMissense | 0.879 | 0.879 | 0.869 | 0.143 | 0.121 | 0.879 | 0.736 |
| VariPred | 0.962 | 0.806 | 0.868 | 0.046 | 0.194 | 0.877 | 0.750 |
| MutPred2 | 0.882 | 0.938 | 0.885 | 0.200 | 0.062 | 0.909 | 0.754 |
| PolyPhen-2 | 0.732 | 0.938 | 0.783 | 0.393 | 0.062 | 0.822 | 0.584 |
| PHACT | **0.048** | 0.031 | 0.164 | 0.690 | 0.969 | 0.038 | -0.692 |
| PHACTboost | 0.821 | 0.958 | 0.882 | 0.185 | 0.042 | 0.885 | 0.775 |
| TrioNsight | 1.000 | 0.913 | **0.944** | **0.000** | 0.087 | **0.955** | **0.890** |

Running TrioNsight with only sequence and structure-based analyses, without the contributions from the other predictors as features, has a detrimental effect on the performance, indicating that the other predictors are important contributors to the high performance. When AlphaMissense, VariPred, PHACT and PHACTboost are all removed from the training the MCC value drops from 0.890 to 0.600 (see Figure 9). This MCC value of 0.600 still outperforms PolyPhen-2, which performs notably lower than the other benchmark methods.

**Figure 9:**
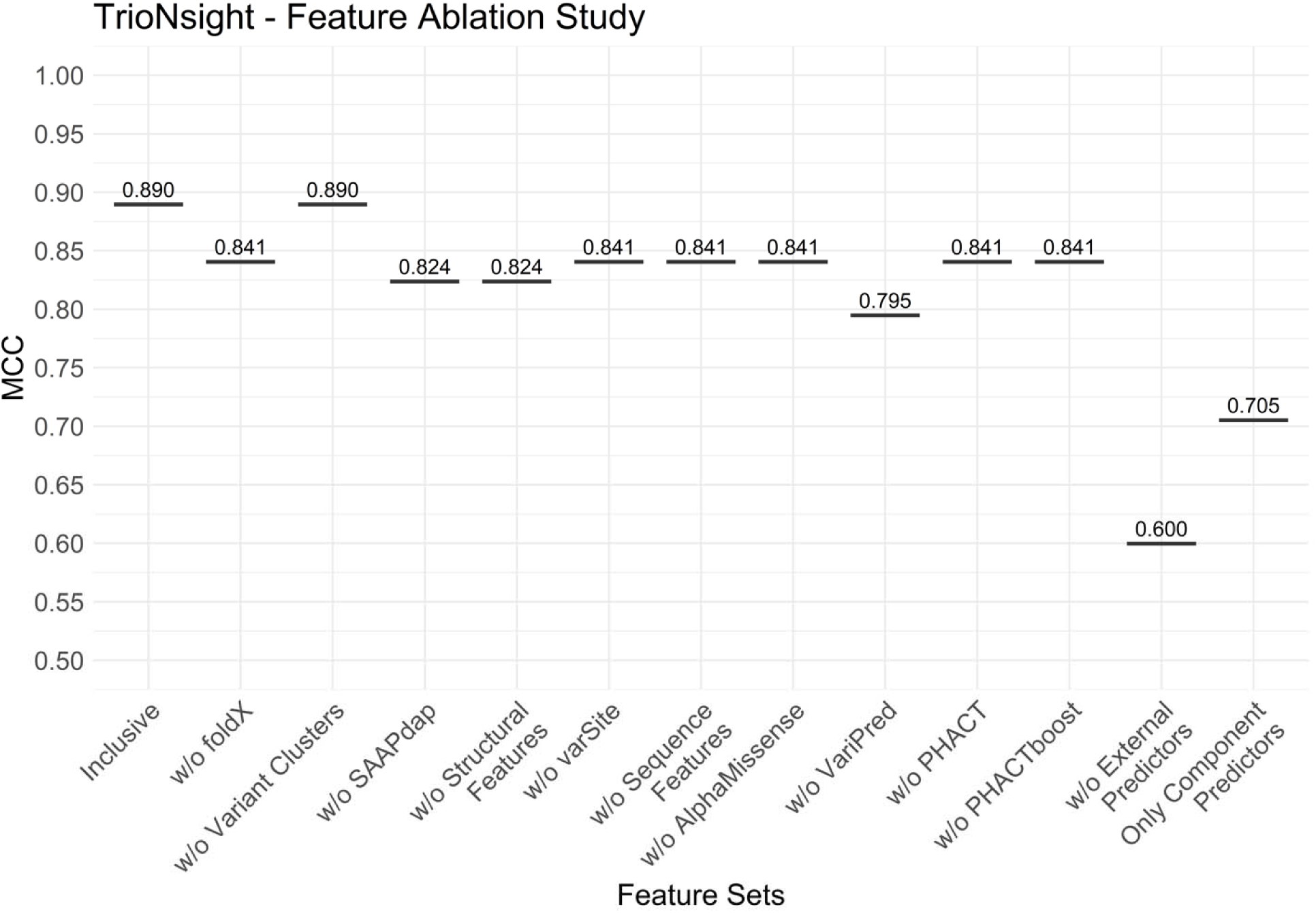
18 different machine learning methods (see Supplementary Table S1) were trained using 13 different feature sets which either use all features in TrioNsight methodology (see Figure 4) or leave one or more features out of the training. ‘Inclusive’ includes all the features, whereas other configurations excluded features as indicated by their labels. Broader exclusions included: ‘w/o Structural Features’ (removing FoldX, Variant Clusters, and all SAAPdap-derived features), ‘w/o Sequence Features’ (excluding all six sequence-based features), and ‘w/o External Predictors’ (omitting AlphaMissense, VariPred, PHACT, and PHACTboost). The ‘Only Component Predictors’ configuration functioned as a meta-predictor using exclusively features from the four external prediction algorithms. A detailed feature ablation analysis demonstrating the contribution of individual features to model performance is provided in Supplementary Table S6.

Individual feature ablation revealed differential contributions amongst the component predictors. VariPred removal had the highest impact, reducing MCC from 0.890 to 0.795. Removing AlphaMissense, PHACT, or PHACTboost individually all resulted in smaller performance decreases, with MCC values dropping to 0.841 in each case. Thus, the structural and sequence-based features developed in this work contributed significantly to enhancing our model’s performance.

TrioNsight was also trained using only features from component predictors as a single meta-predictor, excluding structural and sequence-based features. This approach aimed to measure the contributions of external predictors to TrioNsight’s performance (Figure 9). The MCC value for this model was 0.705, which surpassed the performance of PolyPhen-2, but fell short of all the others.

TrioNsight exhibits robust predictive performance on ambiguous variants, correctly identifying both pathogenic and benign cases that mislead other leading predictors. Table 2 highlights 9 variants that were misclassified by other predictors but correctly identified by TrioNsight. Among these, the variants ARHGEF9 p.R290H and TRIO p.N1465S are noteworthy examples. ARHGEF9 p.R290H weakens the interaction between the DH and PH domains of ARHGEF9 leading to an increased conformational flexibility of the PH domain relative to the DH domain, and is associated with epilepsy and intellectual disability [70–72]. The TRIO p.N1465S mutation impairs TRIO’s ability to activate the small GTPase Rac1, which often leads to neurodevelopmental disorders through decreased neurite outgrowth [73].

**Table 2:** Selected variants within TrioN-like DH domain dataset that are misclassified by other predictors and correctly classified by TrioNsight. The variants are mapped to the corresponding UniProt canonical sequence. Clinical annotation of each variant is supplied in the Annotated column. Correct classifications are bolded for each predictor.

| Protein | Amino acid | Genomic Pos. | Annotated | AlphaMissense | PHACTboost | VariPred | MutPred2 | CADD | ESM-1b | PolyPhen-2 | TrioNsight |
| --- | --- | --- | --- | --- | --- | --- | --- | --- | --- | --- | --- |
| ARHGEF9 | E143K | X 63697259 C T | Benign | **Benign** | Pathogenic | Pathogenic | Pathogenic | **Benign** | Pathogenic | **Benign** | **Benign** |
| ARHGEF9 | R290H | X 63674093 C T | Pathogenic | **Pathogenic** | **Pathogenic** | Benign | **Pathogenic** | **Pathogenic** | **Pathogenic** | **Pathogenic** | **Pathogenic** |
| **TRIO** | H2051Y | 5 14476961 C T | Benign | **Benign** | Pathogenic | **Benign** | Pathogenic | **Benign** | **Benign** | Pathogenic | **Benign** |
| **TRIO** | I1418V | 5 14394071 A G | Benign | **Benign** | Pathogenic | **Benign** | **Benign** | **Benign** | Pathogenic | **Benign** | **Benign** |
| **TRIO** | N1465S | 5 14397125 A G | Pathogenic | **Pathogenic** | **Pathogenic** | Benign | **Pathogenic** | Benign | **Pathogenic** | **Pathogenic** | **Pathogenic** |
| **MCF2** | N594D | X 139602462 T C | Benign | Pathogenic | **Benign** | **Benign** | Pathogenic | Pathogenic | Pathogenic | Pathogenic | **Benign** |
| **PLEKHG2** | F158L | 19 39415432 T C | Benign | Pathogenic | **Benign** | **Benign** | Pathogenic | **Benign** | **Benign** | **Benign** | **Benign** |
| **PLEKHG2** | S226W | 19 39416933 C G | Benign | Pathogenic | Pathogenic | **Benign** | **Benign** | **Benign** | Pathogenic | Pathogenic | **Benign** |
| **PLEKHG1** | A188G | 6 150786440 C G | Benign | Pathogenic | Pathogenic | **Benign** | Pathogenic | Pathogenic | Pathogenic | Pathogenic | **Benign** |

Potential data leakage issues were considered in the training of TrioNsight, and rigorous validation protocols were implemented. (See details in Supplementary Section ‘<u>Data circularity</u> <u>tests</u>’). To assess potential bias, we compared model performance on the initial dataset (with overlapping variants) versus the strict overlap-removed dataset. TrioNsight demonstrated greater robustness to data circularity issues, with a more modest performance increase when overlaps were included compared with the other top-performing methods that experienced greater improvements in their MCC values. When overlaps were removed, TrioNsight maintained its position as the top-performing method with or without overlaps. TrioNsight was able to outperform the benchmark methods in all other metrics, which identifies it as a valuable tool in predicting pathogenicity in the 13 human DH domains contained in the TrioN-like DH domains dataset.

See Supplementary Section ‘Extended performance evaluation of TrioNsight’ for a detailed performance evaluation of TrioNsight model without the exclusion of overlapping variants.

We acknowledge that data circularity might still have caused some inflation in the performance of TrioNsight. However, this effect was minimal and even accounting for this bias, TrioNsight performs better than its competitors, some of which do suffer from data circularity, by a large margin. This robust performance suggests that TrioNsight is a powerful method to predict the variant impacts in TrioN-like DH domains.

Furthermore, these results also demonstrate that using parameters specific to the underlying biology is an effective approach for developing predictors. This strategy allows the incorporation of domain-specific knowledge, leading to more biologically relevant predictions.

### Generalisable workflow for domain-specific variant impact prediction

A standardised workflow that can readily be adapted to create domain-specific variant impact predictors for other protein families has been established by our approach to developing TrioNsight. The methodology employed involves several key steps that could serve as a template for similar efforts across the proteome.

1. A protein domain of interest was identified and classified within functional families to establish evolutionary relationships.
2. A comprehensive dataset was compiled using structural and sequence information from homologous domains across multiple species.
3. Feature engineering integrated domain-specific structural and evolutionary features with predictions from existing tools to incorporate domain-specific context.
4. A meta-predictor was employed that allows for systematic evaluation and selection of the most effective combination of features and prediction methods.

The robust performance of TrioNsight demonstrates that substantial improvements can be achieved through this domain-specific approach in variant impact prediction compared with pan- genome tools. Similar workflows could be particularly valuable for other protein domains associated with disease, such as kinase domains, ion channels, or transcription factor binding domains.

## Conclusions

We present a novel variant impact meta-predictor, TrioNsight, specific to TrioN-like DH domains. TrioNsight deviates from conventional automated variant impact prediction tools, by exploiting both general predictors and DH domain-specific features.

The main difficulty in developing a domain-specific variant impact predictor is the limited amount of variant data available for the training process. By adopting a method to detect functionally similar DH domains with high probability and merging them into a TrioN-like DH dataset, we were able to increase considerably the amount of variant data further enhanced by the variant inheritance method. Consequently, we achieved a larger training dataset that is less prone to overfitting issues.

TrioNsight without the component predictors achieved comparable performance to the six benchmarks. However, the component predictors contribute significantly to the performance of TrioNsight.

A simplified TrioNsight trained using only the component predictor outputs as features, achieved an MCC of 0.705 compared to 0.890 for the full TrioNsight model. Conversely, TrioNsight trained exclusively using features from sequence-based and structural analyses (without the component predictors) achieved an MCC of 0.600. While the component predictors alone yielded competitive results, this analysis clearly demonstrates the value added by our domain-specific feature engineering approach and validates the biological relevance and discriminatory capacity of the TrioN-like DH domain-specific features.

TrioNsight, in the final model, trained using all available features, outperformed all six benchmarking methods in predicting pathogenicity, indicating that selecting biologically relevant features is a successful method for developing a predictor. Therefore, this targeted methodology addresses a significant limitation of existing tools - the over-generalisation of parameters across diverse protein families. Future work could explore the application of this methodology to other protein domains leading to more accurate variant impact prediction tools.

Furthermore, this study provides comprehensive results from both sequence and structure conservation analyses, along with the identification of pathogenic and benign variant clusters on target domains including those that are misclassified by the component predictors. These findings contribute valuable insights into the functional characteristics of highly conserved DH domains. Beyond its specific application to TRIO-related proteins, the systematic workflow developed here - from domain classification to meta-predictor training - establishes a template that can be adapted to create domain-specific variant impact predictors for other protein families, potentially enhancing variant interpretation across the proteome.

## Key points

- TrioNsight is a naïve Bayes meta-predictor designed specifically for variant impact prediction in the N-terminal Dbl-homology domain of TRIO and 12 highly similar human DH domains.
- By focusing on domain-specific biology rather than relying solely on generalised mechanisms, TrioNsight captures features tailored to DH domain function that broader predictors do not exploit.
- The predictor integrates structural, evolutionary, and physiochemical features from approximately 1,500 DH domains across 294 species, providing novel insights into the functional constraints shaping this domain family.
- TrioNsight outperforms all existing general predictors, including AlphaMissense, achieving a Matthews’ Correlation Coefficient of 0.890, and correctly classifies variants that general tools misclassify.
- We provide a comprehensive variant impact map for clinical interpretation and establish a standardised workflow adaptable for developing domain-specific predictors for other protein families.

## Ethics approval statement

This study utilised pre-existing, anonymised data from publicly available and licensed databases (including gnomAD, ClinVar, and HGMD Pro). As the research involved no direct participation of human or animal subjects, institutional ethical approval and informed consent were not required.

## Data availability

The input data, results and code are available at https://github.com/alptaciroglu/TrioNsight

The SAAPdap pipeline is available at https://github.com/ACRMGroup/SAAP3/

Strucclus is available at http://www.bioinf.org.uk/software/strucclus/

## Acknowledgements

We would like to thank Dr. Nicola Bordin for his help in obtaining FunFams assignments of Dbl-like homology domains in the CATH-Gene3D resource base, and for developing the scorecons.py function to run ScoreCons without alignment size limitation, Dr. Ian Sillitoe for his assistance in modelling Dbl-like homology domains together with Pleckstrin homology domains using AlphaFold2, Dr. Paul Ashford for his valuable discussions on the 3D-clustering of variants on protein structures, Dr. Vaishali Waman for her help in generating high-resolution structural models using PyMOL, Weining Lin for her insightful discussions on state-of-the-art variant impact predictors and PI Ogün Adebali for granting access to Sabancı University Troya HPC Cluster.

## References

[1] Schmidt S, Debant A. Function and regulation of the Rho guanine nucleotide exchange factor Trio. Small GTPases 2014;5. 10.4161/sgtp.29769.

[2] Kratzer MC, England L, Apel D, et al. Evolution of the Rho guanine nucleotide exchange factors Kalirin and Trio and their gene expression in Xenopus development. Gene Expression Patterns 2019;32:18–27. 10.1016/J.GEP.2019.02.004.

[3] Blangy A, Vignal E, Schmidt S, et al. TrioGEF1 controls Rac- and Cdc42-dependent cell structures through the direct activation of RhoG. J Cell Sci 2000;113. 10.1242/jcs.113.4.729.

[4] Paskus JD, Herring BE, Roche KW. Kalirin and Trio: RhoGEFs in synaptic transmission, plasticity, and complex brain disorders. Trends Neurosci 2020;43:505. 10.1016/J.TINS.2020.05.002.

[5] Kroon J, Heemskerk N, Kalsbeek MJT, et al. Flow-induced endothelial cell alignment requires the RhoGEF Trio as a scaffold protein to polarize active Rac1 distribution. Mol Biol Cell 2017;28. 10.1091/mbc.E16-06-0389.

[6] Chhatriwala MK, Betts L, Worthylake DK, et al. The DH and PH Domains of Trio Coordinately Engage Rho GTPases for their Efficient Activation. J Mol Biol 2007;368. 10.1016/j.jmb.2007.02.060.

[7] Katrancha SM, Wu Y, Zhu M, et al. Neurodevelopmental disease-associated de novo mutations and rare sequence variants affect TRIO GDP/GTP exchange factor activity. Hum Mol Genet 2017;26. 10.1093/hmg/ddx355.

[8] Genovese G, Fromer M, Stahl EA, et al. Increased burden of ultra-rare protein-altering variants among 4,877 individuals with schizophrenia. Nat Neurosci 2016;19. 10.1038/nn.4402.

[9] Katrancha SM, Shaw JE, Zhao AY, et al. Trio Haploinsufficiency Causes Neurodevelopmental Disease-Associated Deficits. Cell Rep 2019;26. 10.1016/j.celrep.2019.02.022.

[10] Jaiswal M, Dvorsky R, Ahmadian MR. Deciphering the molecular and functional basis of Dbl family proteins: A novel systematic approach toward classification of selective activation of the Rho family proteins. Journal of Biological Chemistry 2013;288. 10.1074/jbc.M112.429746.

[11] Golen K Van. The Rho GTPases in cancer. 2010. 10.1007/978-1-4419-1111-7.

[12] Schmidt A, Hall A. Guanine nucleotide exchange factors for Rho GTPases: Turning on the switch. Genes Dev 2002;16. 10.1101/gad.1003302.

[13] Boland A, Côté JF, Barford D. Structural biology of DOCK-family guanine nucleotide exchange factors. FEBS Lett 2023;597. 10.1002/1873-3468.14523.

[14] Hashim IF, Ahmad Mokhtar AM. Small Rho GTPases and their associated RhoGEFs mutations promote immunological defects in primary immunodeficiencies. International Journal of Biochemistry and Cell Biology 2021;137. 10.1016/j.biocel.2021.106034.

[15] Raimondi F, Felline A, Fanelli F. Catching Functional Modes and Structural Communication in Dbl Family Rho Guanine Nucleotide Exchange Factors. J Chem Inf Model 2015;55. 10.1021/acs.jcim.5b00122.

[16] Toma-Fukai S, Shimizu T. Structural insights into the regulation mechanism of small GTPases by GEFs. Molecules 2019;24. 10.3390/molecules24183308.

[17] Bottani A, Orrico A, Galli L, et al. Unilateral focal polymicrogyria in a patient with classical Aarskog-Scott syndrome due to a novel missense mutation in an evolutionary conserved RhoGEF domain of the faciogenital dysplasia gene FGD1. Am J Med Genet A 2007;143. 10.1002/ajmg.a.31733.

[18] Morimoto S. Sarcomeric proteins and inherited cardiomyopathies. Cardiovasc Res 2008;77:659–66. 10.1093/cvr/cvm084.

[19] Sahai E, Marshall CJ. RHO - GTPases and cancer. Nat Rev Cancer 2002;2. 10.1038/nrc725.

[20] Rao S, Sadybekov A, DeWitt DC, et al. Detection of autism spectrum disorder-related pathogenic trio variants by a novel structure-based approach. Mol Autism 2024;15:12. 10.1186/S13229-024-00590-9.

[21] Von Bohlen Und Halbach O. Dendritic spine abnormalities in mental retardation. Cell Tissue Res 2010;342. 10.1007/s00441-010-1070-9.

[22] Barbosa S, Greville-Heygate S, Bonnet M, et al. Opposite Modulation of RAC1 by Mutations in TRIO Is Associated with Distinct, Domain-Specific Neurodevelopmental Disorders. Am J Hum Genet 2020;106. 10.1016/j.ajhg.2020.01.018.

[23] Skowronek KR, Guo F, Zheng Y, et al. The C-terminal basic tail of RhoG assists the guanine nucleotide exchange factor Trio in binding to phospholipids. Journal of Biological Chemistry 2004;279. 10.1074/jbc.M312677200.

[24] Landrum MJ, Lee JM, Benson M, et al. ClinVar: Improving access to variant interpretations and supporting evidence. Nucleic Acids Res 2018;46. 10.1093/nar/gkx1153.

[25] Karczewski KJ, Francioli LC, Tiao G, et al. The mutational constraint spectrum quantified from variation in 141,456 humans. Nature 2020;581:434–43. 10.1038/s41586-020-2308-7.

[26] Stenson PD, Mort M, Ball E V., et al. The Human Gene Mutation Database (HGMD®): optimizing its use in a clinical diagnostic or research setting. Hum Genet 2020;139. 10.1007/s00439-020-02199-3.

[27] Zaucha J, Heinzinger M, Tarnovskaya S, et al. Family-specific analysis of variant pathogenicity prediction tools. NAR Genom Bioinform 2020;2. 10.1093/nargab/lqaa014.

[28] Bu F, Zhong M, Chen Q, et al. DVPred: a disease-specific prediction tool for variant pathogenicity classification for hearing loss. Hum Genet 2022;141. 10.1007/s00439-022-02440-1.

[29] Jain S, Bakolitsa C, Brenner SE, et al. CAGI, the Critical Assessment of Genome Interpretation, establishes progress and prospects for computational genetic variant interpretation methods. Genome Biology 2024 25:1 2024;25:1–46. 10.1186/S13059-023-03113-6.

[30] Lin W, Wells J, Wang Z, et al. Enhancing missense variant pathogenicity prediction with protein language models using VariPred. Scientific Reports 2024 14:1 2024;14:1–13. 10.1038/s41598-024-51489-7.

[31] Cheng J, Novati G, Pan J, et al. Accurate proteome-wide missense variant effect prediction with AlphaMissense. Science (1979) 2023;381. 10.1126/science.adg7492.

[32] Kuru N, Dereli O, Akkoyun E, et al. PHACT: Phylogeny-Aware Computing of Tolerance for Missense Mutations. Mol Biol Evol 2022;39. 10.1093/molbev/msac114.

[33] Dereli O, Kuru N, Akkoyun E, et al. PHACTboost: A Phylogeny-Aware Pathogenicity Predictor for Missense Mutations via Boosting. Mol Biol Evol 2024;41. 10.1093/MOLBEV/MSAE136.

[34] Brandes N, Goldman G, Wang CH, et al. Genome-wide prediction of disease variant effects with a deep protein language model. Nat Genet 2023;55. 10.1038/s41588-023-01465-0.

[35] Rentzsch P, Witten D, Cooper GM, et al. CADD: predicting the deleteriousness of variants throughout the human genome. Nucleic Acids Res 2019;47:D886–94. 10.1093/NAR/GKY1016.

[36] Pejaver V, Urresti J, Lugo-Martinez J, et al. Inferring the molecular and phenotypic impact of amino acid variants with MutPred2. Nat Commun 2020;11. 10.1038/s41467-020-19669-x.

[37] Adzhubei IA, Schmidt S, Peshkin L, et al. A method and server for predicting damaging missense mutations. Nat Methods 2010;7. 10.1038/nmeth0410-248.

[38] Lewis TE, Sillitoe I, Dawson N, et al. Gene3D: Extensive prediction of globular domains in proteins. Nucleic Acids Res 2018;46. 10.1093/nar/gkx1069.

[39] Sillitoe I, Cuff AL, Dessailly BH, et al. New functional families (FunFams) in CATH to improve the mapping of conserved functional sites to 3D structures. Nucleic Acids Res 2013;41. 10.1093/nar/gks1211.

[40] Jumper J, Evans R, Pritzel A, et al. Highly accurate protein structure prediction with AlphaFold. Nature 2021 596:7873 2021;596:583–9. 10.1038/s41586-021-03819-2.

[41] Consortium TU, Bateman A, Martin M-J, et al. UniProt: the Universal Protein Knowledgebase in 2025. Nucleic Acids Res 2025;53:D609–17. 10.1093/NAR/GKAE1010.

[42] Steinegger M, Meier M, Mirdita M, et al. HH-suite3 for fast remote homology detection and deep protein annotation. BMC Bioinformatics 2019;20. 10.1186/s12859-019-3019-7.

[43] Orengo CA, Taylor WR. SSAP: sequential structure alignment program for protein structure comparison. Methods Enzymol 1996;266. 10.1016/s0076-6879(96)66038-8.

[44] Horne J, Shukla D. Recent Advances in Machine Learning Variant Effect Prediction Tools for Protein Engineering. Ind Eng Chem Res 2022;61. 10.1021/acs.iecr.1c04943.

[45] Vihinen M. How to evaluate performance of prediction methods? Measures and their interpretation in variation effect analysis. BMC Genomics 2012;13 Suppl 4. 10.1186/1471-2164-13-S4-S2.

[46] Peterson TA, Nehrt NL, Park DH, et al. Incorporating molecular and functional context into the analysis and prioritization of human variants associated with cancer. Journal of the American Medical Informatics Association 2012;19. 10.1136/amiajnl-2011-000655.

[47] Miller ML, Reznik E, Gauthier NP, et al. Pan-Cancer Analysis of Mutation Hotspots in Protein Domains. Cell Syst 2015;1. 10.1016/j.cels.2015.08.014.

[48] Ashford P, Pang CSM, Moya-García AA, et al. A CATH domain functional family based approach to identify putative cancer driver genes and driver mutations. Sci Rep 2019;9. 10.1038/S41598-018-36401-4.

[49] Dong R, Peng Z, Zhang Y, et al. MTM-align: An algorithm for fast and accurate multiple protein structure alignment. Bioinformatics 2018;34. 10.1093/bioinformatics/btx828.

[50] Al-Numair NS, Lopes L, Syrris P, et al. The structural effects of mutations can aid in differential phenotype prediction of beta-myosin heavy chain (Myosin-7) missense variants. Bioinformatics 2016;32:2947–55. 10.1093/BIOINFORMATICS/BTW362.

[51] Henikoff S, Henikoff JG. Amino acid substitution matrices from protein blocks. Proc Natl Acad Sci U S A 1992;89. 10.1073/pnas.89.22.10915.

[52] Valdar WSJ. Scoring residue conservation. Proteins 2002;48:227–41. 10.1002/PROT.10146.

[53] Grantham R. Amino Acid Difference Formula to Help Explain Protein Evolution. Science (1979) 1974;185:862–4. 10.1126/SCIENCE.185.4154.862.

[54] Al-Numair NS, Martin ACR. The SAAP pipeline and database: tools to analyze the impact and predict the pathogenicity of mutations. BMC Genomics 2013;14:S4. 10.1186/1471-2164-14-S3-S4.

[55] Laskowski RA, Stephenson JD, Sillitoe I, et al. VarSite: Disease variants and protein structure. Protein Sci 2020;29:111–9. 10.1002/PRO.3746.

[56] Stephenson JD, Totoo P, Burke DF, et al. ProtVar: mapping and contextualizing human missense variation. Nucleic Acids Res 2024;52:W140–7. 10.1093/NAR/GKAE413.

[57] Delgado J, Radusky LG, Cianferoni D, et al. FoldX 5.0: working with RNA, small molecules and a new graphical interface. Bioinformatics 2019;35:4168–9. 10.1093/BIOINFORMATICS/BTZ184.

[58] Kuhn M. Classification and Regression Training [R package caret version 7.0-1]. CRAN: Contributed Packages 2024. 10.32614/CRAN.PACKAGE.CARET.

[59] Fort P, Blangy A. The evolutionary landscape of Dbl-like RhoGEF families: Adapting eukaryotic cells to environmental signals. Genome Biol Evol 2017;9. 10.1093/gbe/evx100.

[60] Loirand G, Scalbert E, Bril A, et al. Rho exchange factors in the cardiovascular system. Curr Opin Pharmacol 2008;8. 10.1016/j.coph.2007.12.006.

[61] Nile AH, Bankaitis VA, Grabon A. Mammalian diseases of phosphatidylinositol transfer proteins and their homologs. Clin Lipidol 2010;5. 10.2217/clp.10.67.

[62] Momotani K, Somlyo A V. P63RhoGEF: A New Switch for Gq-Mediated Activation of Smooth Muscle. Trends Cardiovasc Med 2012;22. 10.1016/j.tcm.2012.07.007.

[63] Liu X, Wang H, Eberstadt M, et al. NMR structure and mutagenesis of the N-terminal Dbl homology domain of the nucleotide exchange factor Trio. Cell 1998;95. 10.1016/S0092-8674(00)81757-2.

[64] Komai K, Mukae-Sakairi N, Kitagawa M, et al. Characterization of novel splicing variants of the mouse MCF-2 (DBL) proto-oncogene. Biochem Biophys Res Commun 2003;309. 10.1016/j.bbrc.2003.08.088.

[65] Wu JH, Fanaroff AC, Sharma KC, et al. Kalirin promotes neointimal hyperplasia by activating rac in smooth muscle cells. Arterioscler Thromb Vasc Biol 2013;33:702–8. 10.1161/ATVBAHA.112.300234.

[66] Maiwald S, Motazacker MM, Van Capelleveen JC, et al. A rare variant in MCF2L identified using exclusion linkage in a pedigree with premature atherosclerosis. European Journal of Human Genetics 2016;24. 10.1038/ejhg.2015.70.

[67] Chicco D, Jurman G. The advantages of the Matthews correlation coefficient (MCC) over F1 score and accuracy in binary classification evaluation. BMC Genomics 2020;21. 10.1186/s12864-019-6413-7.

[68] Lobo JM, Jiménez-valverde A, Real R. AUC: A misleading measure of the performance of predictive distribution models. Global Ecology and Biogeography 2008;17. 10.1111/j.1466-8238.2007.00358.x.

[69] Chicco D, Jurman G. The Matthews correlation coefficient (MCC) should replace the ROC AUC as the standard metric for assessing binary classification. BioData Min 2023;16. 10.1186/s13040-023-00322-4.

[70] Lemke JR, Riesch E, Scheurenbrand T, et al. Targeted next generation sequencing as a diagnostic tool in epileptic disorders. Epilepsia 2012;53. 10.1111/j.1528-1167.2012.03516.x.

[71] Papadopoulos T, Schemm R, Grubmüller H, et al. Lipid binding defects and perturbed synaptogenic activity of a collybistin R290H mutant that causes epilepsy and intellectual disability. Journal of Biological Chemistry 2015;290. 10.1074/jbc.M114.633024.

[72] Long P, May MM, James VM, et al. Missense mutation R338W in ARHGEF9 in a family with X-linked intellectual disability with variable macrocephaly and macro-orchidism. Front Mol Neurosci 2016;8. 10.3389/fnmol.2015.00083.

[73] Gazdagh G, Hunt D, Gonzalez AMC, et al. Extending the phenotypes associated with TRIO gene variants in a cohort of 25 patients and review of the literature. Am J Med Genet A 2023;191. 10.1002/ajmg.a.63194.

