## Supplementary Material Text for "TrioNsight: Building a meta-predictor to evaluate the clinical impact of TrioN-like Dbl-homology domain variants"

### **Supplementary Materials**

### **Component predictors of TrioNsight**

The component predictors used in TrioNsight are: VariPred [1], AlphaMissense [2], PHACT [3] and PHACTboost [4].

VariPred is a supervised deep learning-based classifier powered by protein language models. It does not require a multiple sequence alignment to run, reducing the computational cost of using the algorithm. Although it requires only sequence data, VariPred was shown to outperform its benchmark competitors in most cases, including high-performance predictors such as 3Cnet [5] and REVEL [6]. In CAGI-6, 3Cnet was selected as the top-ranked method in the SickKids clinical genomes and transcriptomes panel and REVEL was the best ensemble model in correctly identifying pathogenic variants.

AlphaMissense is an unsupervised deep learning model that also leverages protein language models. It uses structural information from the well-established AlphaFold2 algorithm together with sequence information. The performance of AlphaMissense was evaluated in multiple benchmarks including ClinVar missense variants, variants in 612 disease-associated genes, and proteins encoded by a set of clinically important genes selected by the American College of Medical Genetics [7] and shown to outperform all benchmark tools against which it was assessed including EVE, REVEL, CADD and ESM, achieving state-of-the-art performance. AlphaMissense pathogenicity predictions have been used in other studies [8,9].

PHACTboost is a gradient-boosting tree-based classifier that uses phylogenetic trees from its predecessor model, PHACT, and combines this with data obtained from reconstructed ancestral sequences and multiple sequence alignments. PHACTboost was shown to achieve either comparable performance, or to outperform all benchmark tools used in its performance evaluation, including AlphaMissense, EVE [10], and CPT-1 [11].

### **Extended methodology for the selection of TrioN-like DH domains**

Our selection protocol comprised a sequence-based analysis using functional families generated by the FunFams algorithm [12], and a structure-based analysis using experimental and AlphaFold2-generated [13] models as described below.

### 1) Sequence-based analysis

DH domain protein sequences were extracted for CATH Superfamily 1.20.900.10 from the CATH-Gene3D database [14]. The list of DH domains containing genes from all organisms was manually curated to select only one transcript of each gene, favouring a higher UniProt annotation score [15] and longer transcript length. At this point, the list of DH domains contained more than 10000 sequences. Although they are assigned to the same CATH Superfamily, some of these domains could be functionally diverse, coming from orthologs and paralogs in a diverse range of organisms across a broad taxonomic spectrum [16].

To select DH domains that are functionally similar to TrioN, protein domain sequences were annotated using the CATH FunFams classification. FunFams are subfamilies within CATH evolutionary superfamilies that contain domains likely to be highly structurally and functionally similar [12]. To merge related FunFams likely to have similar functions, the seed alignments of functional families were compared using the HHAlign function from hhSuite [17] with the e-value threshold of ${10}^{-24}$ (see Supplementary Figure S6 and Supplementary Figure S7).

### 2) Structure-based analysis

For those FunFams with high sequence similarity measured by HHalign, the structural similarity of domains between the FunFams was checked. This approach leverages the well-established association between protein structure and function [18,19]. The FunFams which were found to be similar by HHalign and whose domains were structurally similar by CATH-SSAP [20] (i.e. having an RMSD score < 2.5 Å) were merged to give an expanded data set with more sequences to provide more training data for the predictor.

Structural models were created using AlphaFold2 which provides the pLDDT score as an indicator of model confidence, where scores above 70 indicate confidence in the model. All the models had a pLDDT score ≥ 70 with a mean of 91.06. Using the TrioN DH domain as the reference structure, domain models with high structural similarity to the TrioN DH domain were selected based on a threshold of 2.5 Å Root Mean Square Deviation (RMSD) on Cα atoms. Additionally, a minimum overlap criterion of 80% of residues superposed was used to ensure a significant portion of the domain structures are substantially similar.

The sequence and structure-based analyses yielded a dataset of 1526 DH domains from diverse species which are highly similar in their structure and likely to have functional similarity. The dataset containing these 1526 DH domains is termed the TrioN-like DH dataset. 13 of these are human domains that are assigned to 5 functional families. In addition, AlphaFold2 structure models of human DH domains within the TrioN-like DH dataset were superimposed and visualised to verify their high structural similarity (see Supplementary Figure S7 for the process diagram of the TrioN-like DH dataset creation).

### **Extended methodology for the conservation analysis and variant mapping of TrioN-like DH domains**

Conservation analysis was performed on three DH domain datasets: a representative sample of the entire DH superfamily, the specific FunFam containing TrioN, and the TrioN-like DH domain dataset. The sequences of DH domains in these datasets were multiply aligned using MUSCLE [21] and conservation scores for each position were obtained using the Scorecons method [22]. These scores were used to identify novel functional sites and compare them with known catalytic residues and substrate-binding regions (e.g., Rac1 and RhoG binding regions [23,24]). Scorecons also measures the information content of the alignment, which is determined by the Diversity of Positions (DOPS) score. The DOPS score ranges from 0 to 100, and a DOPS score ≥ 70 is associated with high information content. A DOPS score of 85.1 was obtained for the TrioN-like DH dataset alignment.

Since CATH clusters all the DH domains into one evolutionary superfamily and then subclassifies functionally close relatives into FunFams, we wanted to check that our selection protocol had only merged FunFams that were similar in their functions. If this were not the case, merging the FunFams would remove the conservation signal of functional residues. We therefore compared the conserved residues obtained using:

(1) A representative sample of the whole DH superfamily (10,250 sequences reduced to 1245 sequences). The sample was selected to reduce the computational time by running the MMSeqs2 algorithm with 70% sequence similarity and 80% sequence coverage parameters.

(2) The specific FunFam associated with the N-terminal TRIO DH domain (FF-00001). Both TrioN and KalrnN (the N-terminal DH domain of KALRN) are classified in this functional family, which contained 598 sequences in total.

(3) The TrioN-like DH dataset obtained by expanding the TrioN FunFam with other FunFams whose domains have high sequence and structure similarity with TrioN. This contained 1526 sequences.

All the MSAs contained the representative N-terminal TRIO DH domain (TrioN). Scorecons conservation scores were calculated for all three multiple sequence alignments. The resulting scores were mapped to the TrioN-like DH domain dataset representative and used to colour the structural model to visualise functionally important regions. A blue-white-red colour spectrum, representing scores ranging from 0 (blue) to 100 (red), was used for colouring, and visualised using PyMOL [25].

Following the conservation analysis, nonsynonymous variants were collected for the TrioN-like proteins in our dataset from ClinVar (version 2024/04/01), GnomAD (version 4.1.0 - accessed 2024/05/02), and HGMD Pro (version 2024.2) databases. The Clinical Significance label from both ClinVar and GnomAD was used to select variants from three categories: 1) pathogenic (includes ‘likely pathogenic’ and ‘pathogenic’) 2) benign (includes ‘likely benign’ and ‘benign’) or 3) unknown significance. For the GnomAD database, which was used almost exclusively to obtain benign variants, an allele count ≥ 5 and a minor allele frequency (MAF) of ≥ 0.001 were used as filters to select benign variants. A MAF threshold of ≥ 0.001 has been used successfully in previous genomic studies [26,27].

Variants that mapped onto the corresponding DH domains of the proteins, based on the CATH-Gene3D domain boundaries, were extracted. Variants that map to any other transcript than that of the UniProt canonical sequence (designated with the Ensembl Transcript ID) were discarded, as these might have different functional consequences. After variant collection, a total of 29 benign and 32 pathogenic variants were obtained from 13 human DH domains.

### **Creating AlphaFold models for the TrioN-like DH domains**

Multiple structural models of the TrioN-like DH domains and their surrounding regions were generated using AlphaFold2 and subsequently used in variant inheritance and structural analyses. Owing to the size limitations of AlphaFold2, full models for the larger proteins TRIO and KALRN could not be created. Instead, several partial models were produced for each protein, and the reliable ones were selected based on their pLDDT values (pLDDT score ≥ 70), which indicate model confidence.

### **Feature generation protocols**

### 1) Sequence-based analyses

Three of the 6 sequence-based features were obtained from running the corresponding algorithms on the alignment of the TrioN-like DH cluster sequences: conservation was obtained from the Scorecons algorithm [22]; the BLOSUM80 matrix was used to obtain the likelihood of amino acid substitutions corresponding to variants in the TrioN-like DH dataset; Grantham distance was used to reflect the physicochemical changes in the mutated residues. Functional relevance from UniProtKB/SwissProt is separated into 2 features, a Boolean value to indicate whether a residue is annotated as a feature and a binary field to indicate residue modifications including phosphorylation, methylation and acetylation. Finally, the VarSite conservation scores were retrieved from ProtVar, which are Scorecons scores obtained from UniRef90 sequence alignments separately for each variant.

### 2) Structural analyses

Apart from the Distance to Variant Clusters (DVC) and FoldX, all 11 structural analyses were performed by deploying the SAAPdap pipeline. A typical SAAPdap run analyses a specified nonsynonymous variant for 14 likely detrimental changes. 13 of these changes are impacts on structures likely to be harmful [28–30] as shown in ‘Structural Analyses’ (see Figure 4). Any features that are identical for all variants were removed.

DVC is a metric created to measure the distance of a given position to the variant clusters. Using strucclus, 5 clusters was selected as the most informative clustering threshold based on the p-value and analysis of the distribution of the clusters (see Results for details). To use the variant cluster information to train TrioNsight, the centroids of each of the 5 clusters were calculated and the Cα distance of each variant to each cluster centroid was measured. The DVC comprises 5 features, one for each of the 5 clusters.

ProtVar [31] contains over 200M precalculated stability changes from FoldX v5.0, which are represented in the foldXDdg and foldX_pLDDT columns. The stability changes provided in ProtVar associated with the variant dataset in our analysis were extracted and used in training TrioNsight.

### 3) Variant Prediction Algorithms

The following methods were used to obtain the relevant predictions for each algorithm and the scores corresponding to our variant dataset were extracted. PHACT and PHACTboost prediction scores were downloaded from the link the authors provided under the ‘Data Availability’ section of their study [4]. The ProtVar API wrapper script was used to obtain the AlphaMissense predictions. VariPred was installed and run following the authors' instructions. VariPred variant predictions were obtained without retraining the algorithm.

### **Model selection**

The performance of 18 machine learning methods was evaluated to select the method to use in TrioNsight. The best-performing method was selected using a 2-step test. First, the performances were visualised in terms of MCC values by training 18 methods 20 times each (see Supplementary Figure S8) using all of the TrioNsight features and grouping MCC values by the method. Also, the variance in the MCC values was measured in this step.

Second, the contributions of distinctive features to the performance of the predictor were tested. 18 methods were trained 13 times using different feature configuration in each iteration and performance was grouped using the configuration (see Supplementary Figure S9). These configurations either include all features in the training (Inclusive) or leave at least one (foldX, AlphaMissense etc.) or more features (w/o External Predictors) out of the training. The naïve-Bayes method was selected as the best-performing method.

### **Equations**

- Precision measures the proportion of correctly identified positive predictions among all positive predictions.

$$Precision= \frac{TP}{TP + FP}$$

- Recall quantifies the proportion of actual positive cases that were correctly identified.

$$Recall= \frac{TP}{TP + FN}$$

- The overall Accuracy metric assesses the model's general performance across all predictions.

$$Accuracy= \frac{TP + TN}{TP + TN + FP + FN}$$

- False Positive Rate is an error rate that indicates the proportion of negative cases incorrectly classified as positive.

$$False Positive Rate= \frac{FP}{FP + TN}$$

- False Negative Rate is an error rate that indicates the proportion of positive cases incorrectly classified as negative.

$$False Negative Rate= \frac{FN}{FN + TP}$$

- The F1 Score represents the harmonic mean of precision and recall, which was used to provide a single score that balances both metrics.

$$F1 Score= \frac{2 * Precision * Recall}{Recall + Precision}$$

- The MCC considers all four confusion matrix categories and provides a balanced measure ranging from -1 to +1, where +1 represents perfect prediction, 0 represents random prediction, and -1 represents complete disagreement between prediction and observation.

$$MCC= \frac{TP*TN-FP*FN}{\sqrt{\left( TP+FP \right)\left( TP+FN \right)\left( TN+FP \right)\left( TN+FN \right)}}$$

### **Data circularity tests**

Data circularity is a well-known challenge in variant impact prediction, which arises when the same data are used both to train and evaluate the performance of the predictor [32,33]. To ensure the robustness of the predictive model and minimise the risk of data leakage or circular reasoning, several validation approaches were performed.

It is important to consider the potential overlap between the evaluation dataset of TrioNsight and the training datasets of both its component predictors and the benchmarking predictors. Such data overlap could lead to biased performance evaluations for component predictors and feed-forward into the meta-predictor (TrioNsight). Upon investigation, data overlaps were found in the case of MutPred2, PHACTboost, and VariPred, which had 9, 10, and 8 variants respectively common between their training data and the 61 variants used in the TrioNsight evaluation dataset. Conversely, CADD did not have any training dataset overlap, and PolyPhen-2 had only 1 overlap out of 61 variants. Two predictors were exempt from overlap concerns: AlphaMissense, which uses weak labels in its training process, and PHACT, which employs a probabilistic, phylogeny-dependent scoring method rather than a trained ML model. In total, 25 out of 61 variants in the TrioN DH dataset overlapped in the training set of at least one predictor.

To address these potential biases, benchmarking was carried out excluding overlapping variants for all methods except CADD, ESM-1b, PHACT, and AlphaMissense. For the rest of the methods, overlapping data were removed separately for each method to observe the change in their predictive power. TrioNsight was trained using a leave-one-out cross-validation (LOOCV) approach where cross-validation runs were skipped if the test variant overlapped with component predictor training data. In this modified protocol, training used all variants except the held-out test variant, preventing artificial inflation of performance metrics through data leakage. While it is often the case that general predictors also avoid overlap at the same-protein or same-homologous-family level, such considerations make little sense for a restricted close-family predictor like TrioNsight.

However, as a further assessment of potential bias owing to data circularity, we also compared model performance using the initial dataset of 61 variants, in which variants overlapping with the training sets of component predictors were not excluded. This comparison provides insight into the potential impact of information leakage, as the overlap-removed dataset represents a more stringent testing scenario. The modest difference in MCC scores (0.906 for the initial dataset with overlaps vs. 0.890 for the overlap-removed dataset) suggests that any inflation in performance owing to data circularity was limited. In the analysis including overlapping variants, MutPred2 demonstrated slightly worse MCC values whereas PolyPhen-2, PHACTboost, VariPred and TrioNsight demonstrated slightly better MCC values.

The predictive power of TrioNsight and other component methods remained largely stable in subsequent performance evaluation, indicating that the approach does not rely on circular reasoning from pre-existing methods (see Table 1 for performance evaluation against eight established algorithms using non-overlapping data).

Beyond the leave-one-out cross-validation methodology that was primarily employed throughout this study, an additional validation was performed by training TrioNsight on a subset of the 61 variants and then evaluating its performance on the remaining held-out variants. This validation approach yielded an MCC value of 0.866 for the 14 test variants, which were selected to ensure they did not overlap with component predictor training data, indicating robust classification performance. The strong performance provides additional evidence for the reliability of the predictive framework underlying TrioNsight, complementing the cross-validation results.

Finally, the training dataset was expanded by inheriting the variant collection across multiple structural models. The effective training sample size was substantially increased by this diversification strategy, whilst the risk of overfitting to specific structural contexts was simultaneously reduced.

### **Optimizing strucclus parameters**

Strucclus was run to identify ‘n’ clusters where n varied from 2 to 8. Supplementary Table S5 shows the results as p-values and ꭓ^2^ values. While n=3 gave the best p-value, n=5 ( which gave a slightly worse p-value), was chosen because this segregates residues involved in binding TRIO-Rac1 (see Figure 6) that are enriched in pathogenic variants (see Figure 8). See Supplementary Movie S2 for rotational visualisation of variant clusters on the representative 1NTY TrioN-DH model.

### **Extended performance evaluation of TrioNsight**

Another version of TrioNsight was trained and evaluated to assess the impact of data circularity arising from variants that overlap with the training data of component predictors. Unlike previous analyses that excluded these overlapping variants, this implementation deliberately retains the complete variant dataset when evaluating both TrioNsight and the individual component predictors, enabling direct measurement of how data circularity affects performance across all benchmarking methods.

The performance of TrioNsight trained using the naïve-Bayes algorithm was compared against CADD, ESM-1b, AlphaMissense, VariPred, MutPred2, PolyPhen-2, PHACT and PHACTboost predictors when evaluated on the complete variant dataset, including overlapping variants (see Supplementary Table 7).

Under these conditions with potential data circularity, TrioNsight outperforms all benchmark methods by a wide margin based on the MCC metric (0.906). According to MCC, PHACTboost, VariPred and AlphaMissense are TrioNsight’s closest competitors in that order (MCC = 0.809, 0.782 and 0.736 respectively). PHACTboost and ESM-1b perform better than TrioNsight in Recall and False Negative Rate metrics, which shows their ability to predict pathogenic variants correctly, at the expense of identifying benign variants as pathogenic.

Compared to the benchmarking carried out by removing the overlapping variants (see Table 1), the changes in the MCC values clearly showed that these predictors do not have 100% accuracy on the variants they have encountered during their training. Considering why the change in MCC values was not greater than observed: firstly, the number of overlapping variants was too small for each method; secondly, removed variants resulted in a coincidental compensation where some predictions got better and some worse; finally, these predictors might have been developed to prevent overfitting and perform similarly on seen and unseen data. **This relative stability in TrioNsight's performance when overlapping variants were removed further supports the robustness of our method against data leakage concerns.**

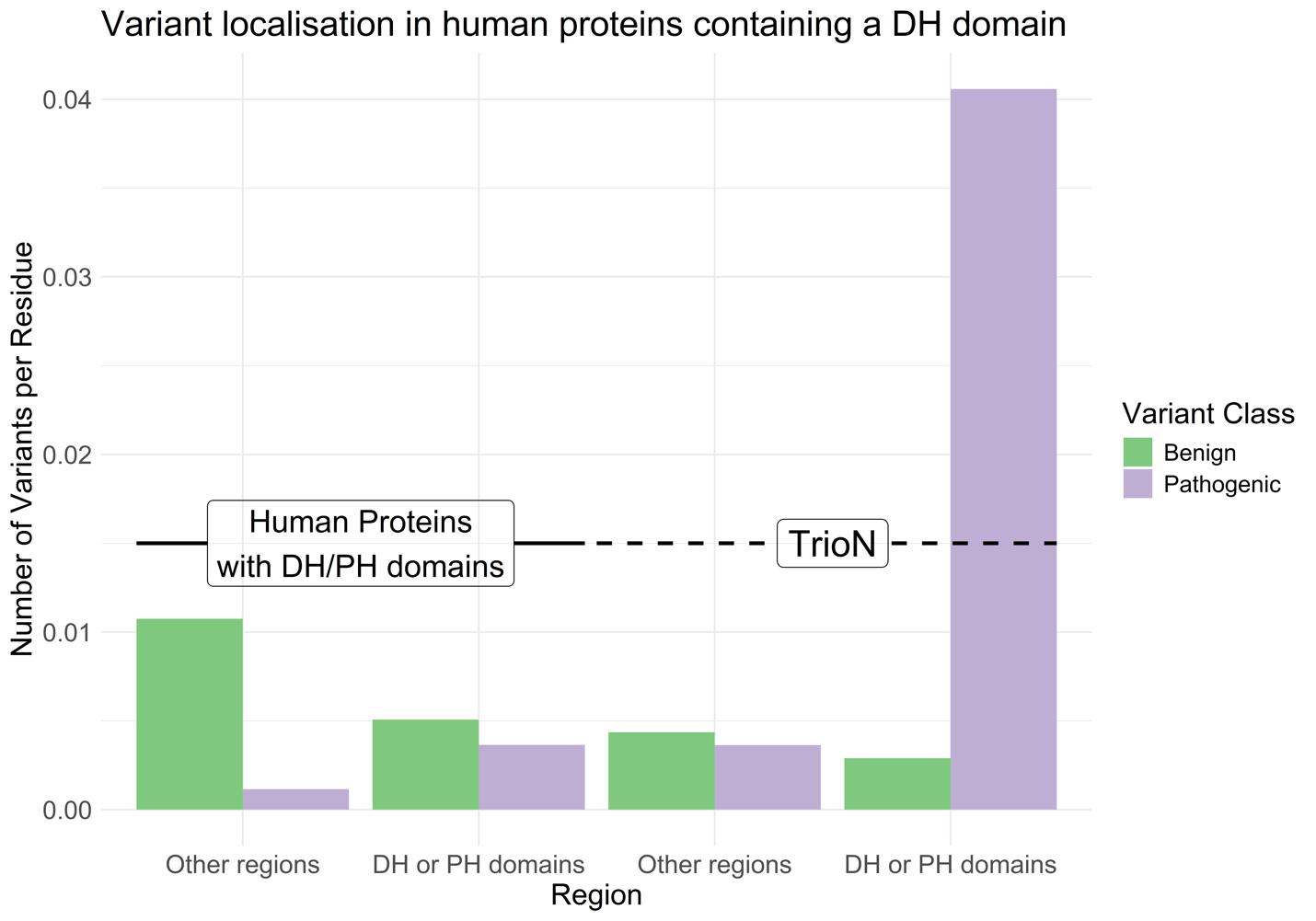
**Supplementary Figure S1:** The distribution of benign and pathogenic variants across human DH domain-containing proteins. The distribution of variants from all 70 human Dbl-homology domains was plotted. The number of variants was divided by the domain length in which the variants were located, resulting in a normalised ratio. N-terminal TRIO DH-PH domain contains a much higher ratio compared to other DH domains.

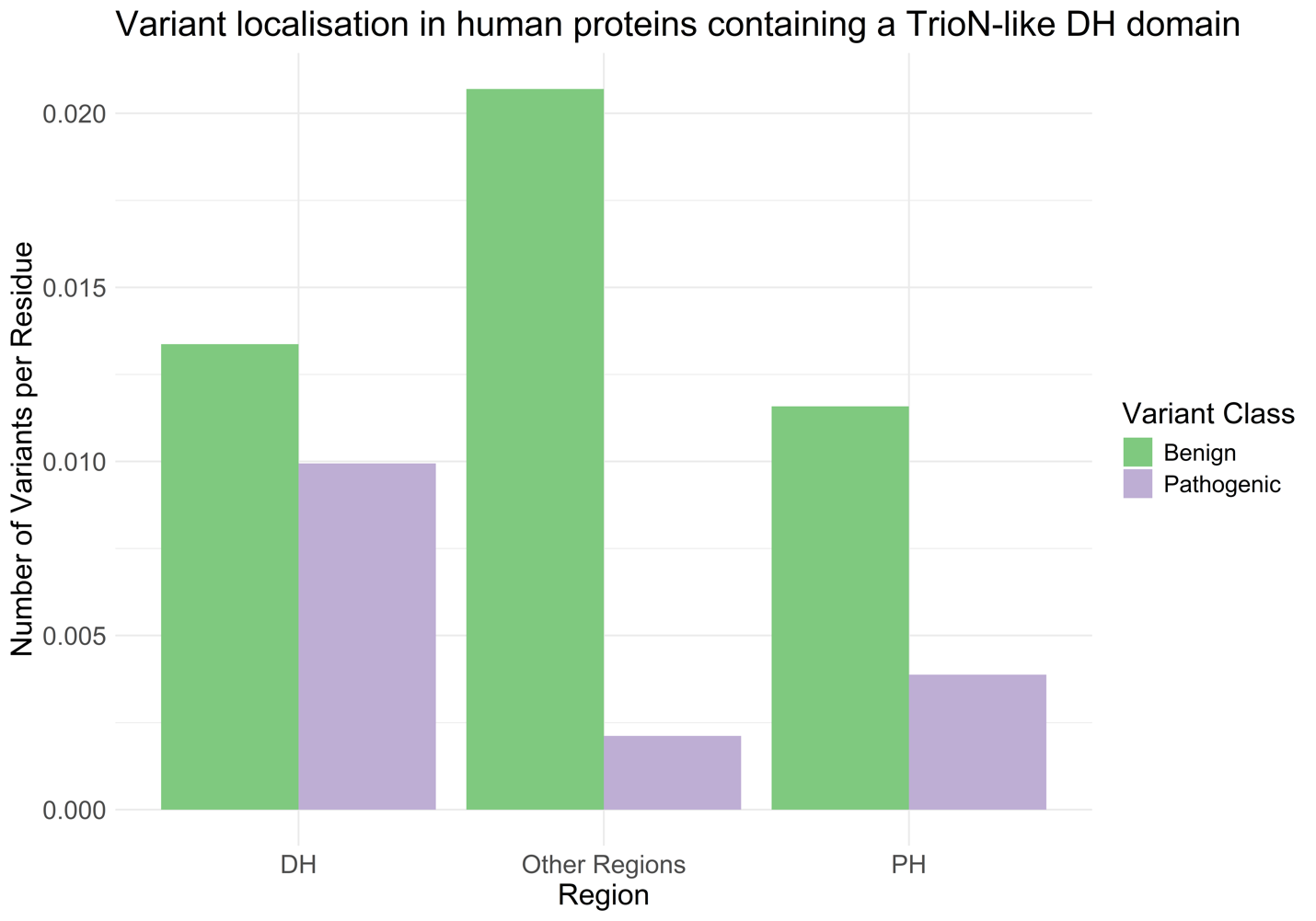

**Supplementary Figure S2:** The distribution of benign and pathogenic variants across proteins containing a TrioN-like DH domain. The distribution of variants from all 13 TrioN-like Dbl-homology domains was plotted. All TrioN-like Dbl-homology domains are from human proteins. The number of variants was divided by the length of the region in which the variants were located, resulting in a normalised ratio. Length of the DH domains, PH domains and whole protein length excluding the DH and PH domains were used to normalise the data

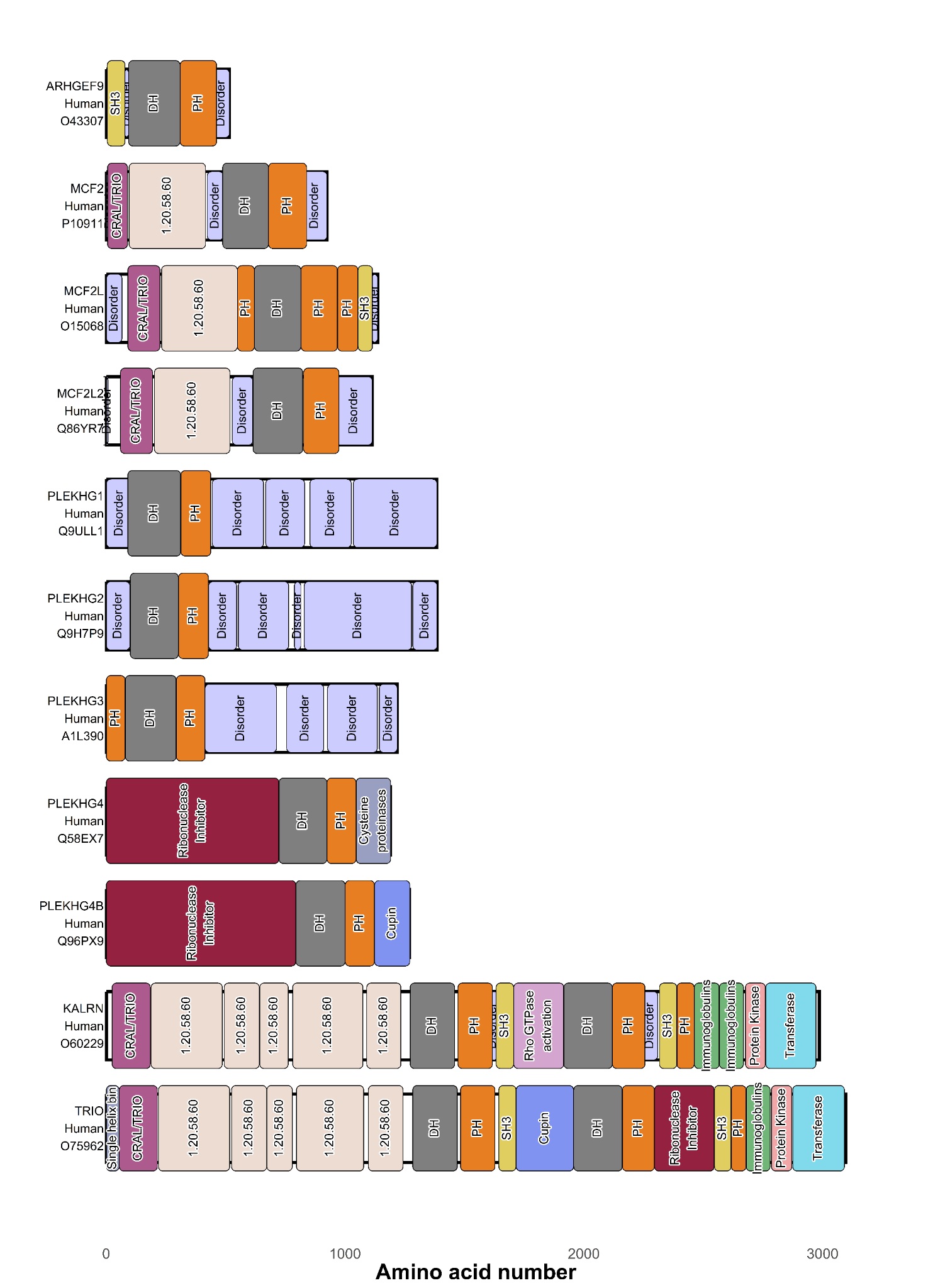

***Supplementary Figure S3:*** Multidomain organisation of human proteins that contain a TrioN-like DH domain. Domain boundaries were obtained from the CATH-Gene3D database. The disordered regions were extracted from the MobiDB database. TRIO and its paralog protein KALRN contain two sets of DH domains named N-terminal or C-terminal DH domains based on the proximity to the termini of the polypeptide.

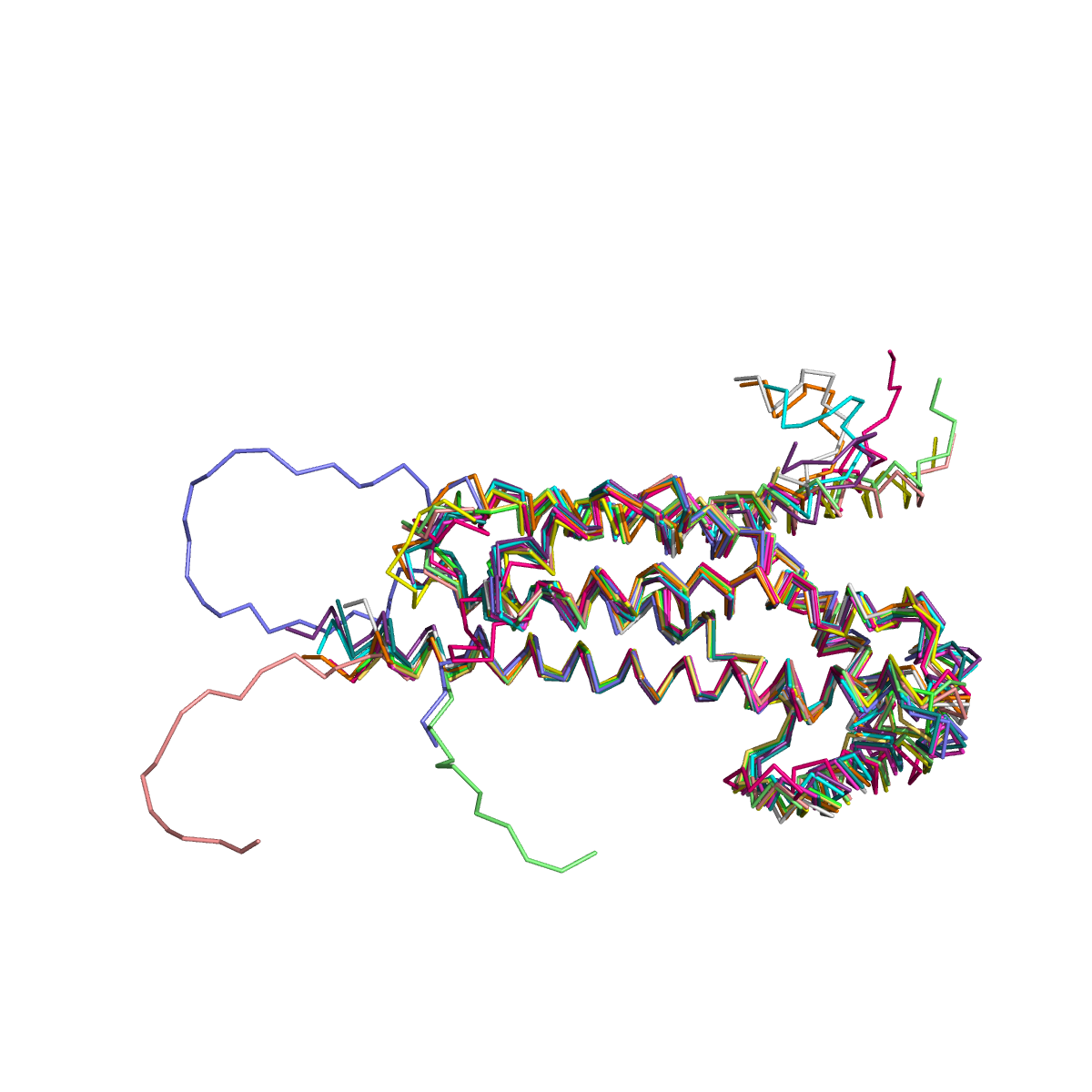

**Supplementary Figure S4:** The superimposition of 13 human DH domains in TrioN-like DH models. The models were created using AlphaFold2. The models were visualised in PyMOL software using ribbon representation. TrioN-like DH domains superimpose very closely, suggesting functional and evolutionary relationships.

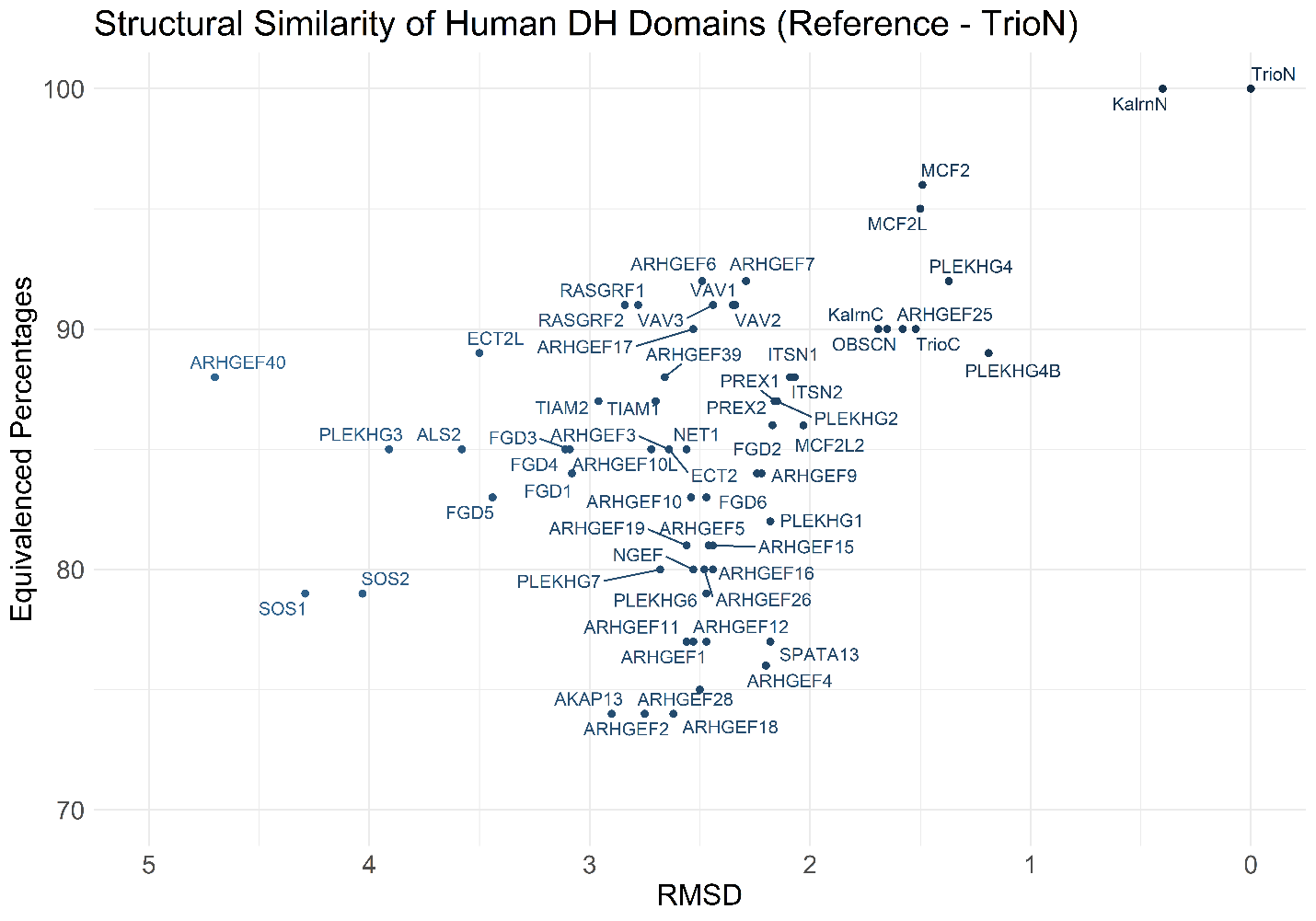

**Supplementary Figure S5:** Root mean square deviation (RMSD) values plotted against equivalenced percentages to visualise the structural similarity between human DH domains. The equivalenced percentages are the length of the aligned regions divided by the length of the longer sequence out of the query structure and reference structure. All human DH domains (query structures) are pairwise aligned to the TrioN (reference structure). RMSD values show how similar those aligned regions are. Most human DH domains are highly similar in terms of structures visualised in the plot.

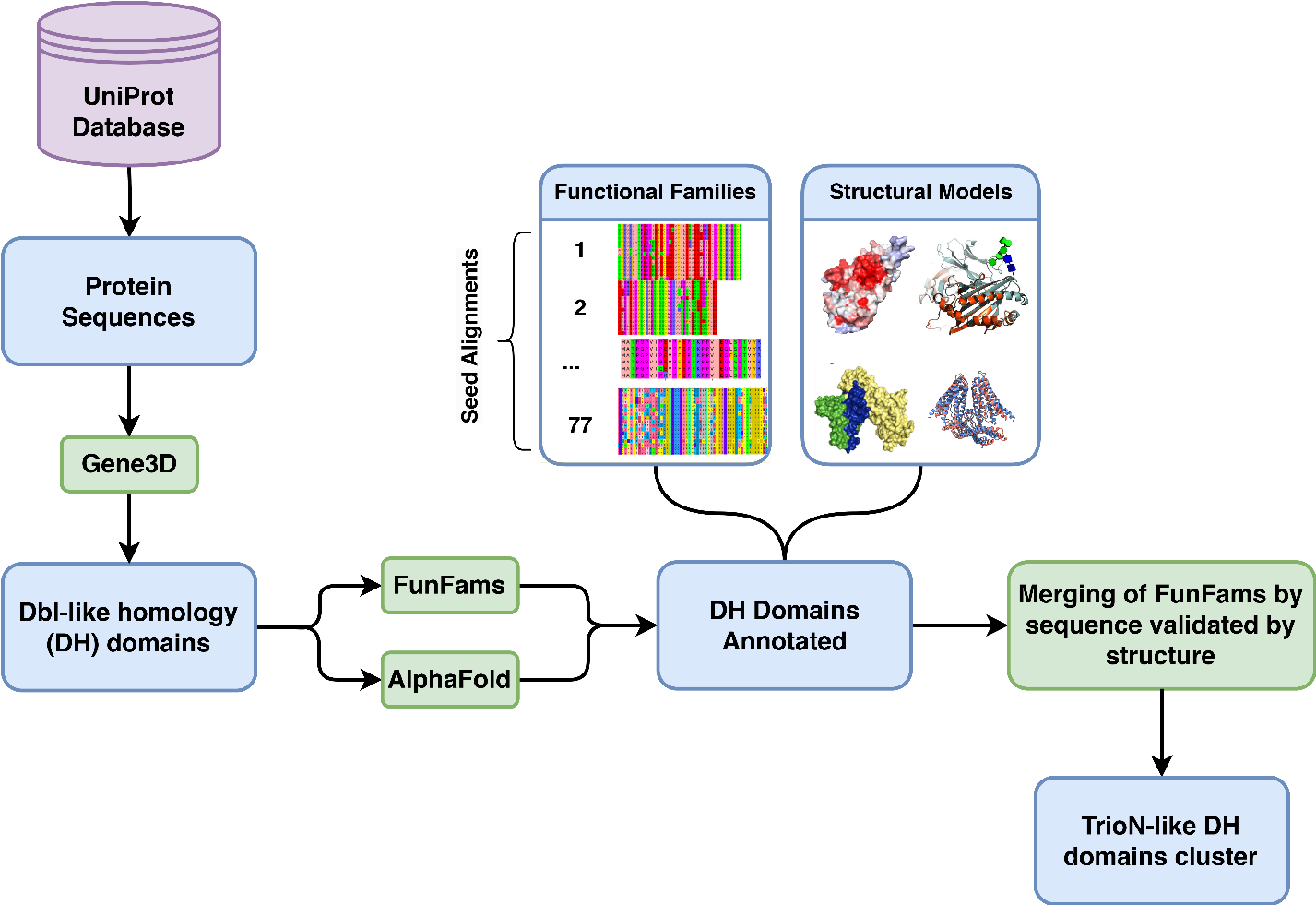

**Supplementary Figure S6:** Workflow to obtain TrioN-like Dbl-homology (DH) domains for TrioNsight. DH domains were identified using CATH-Gene3D [14] domain annotations of the protein sequences from UniProtKB [15]. Structural models were created for each of these DH domains using AlphaFold2. DH domains were assigned to their corresponding functional families using the FunFams algorithm [12]. Subsequently, HHalign [17] was used to identify highly similar FunFams using an e-value threshold of 10^-24^ to identify a larger set of sequences containing the TrioN DH domain. Pairs of FunFams were merged if the relatives were structurally similar. This was determined using structure-based analyses, namely the CATH-SSAP [20] structure comparison method with thresholds of 80% residue overlap and RMSD < 2.5 Å superposition against the reference structure of the TrioN DH domain (see text). This gave a broad set of sequences that contained the TrioN DH domain, but in which relatives were structurally highly similar and likely to be functionally similar. The final subcluster containing TrioN was named the ‘TrioN-like DH domains cluster’. It contains DH domains from FF-000001, FF-000002, FF-000008, FF-000019, and FF-000028 functional families of the CATH superfamily 1.20.900.10.

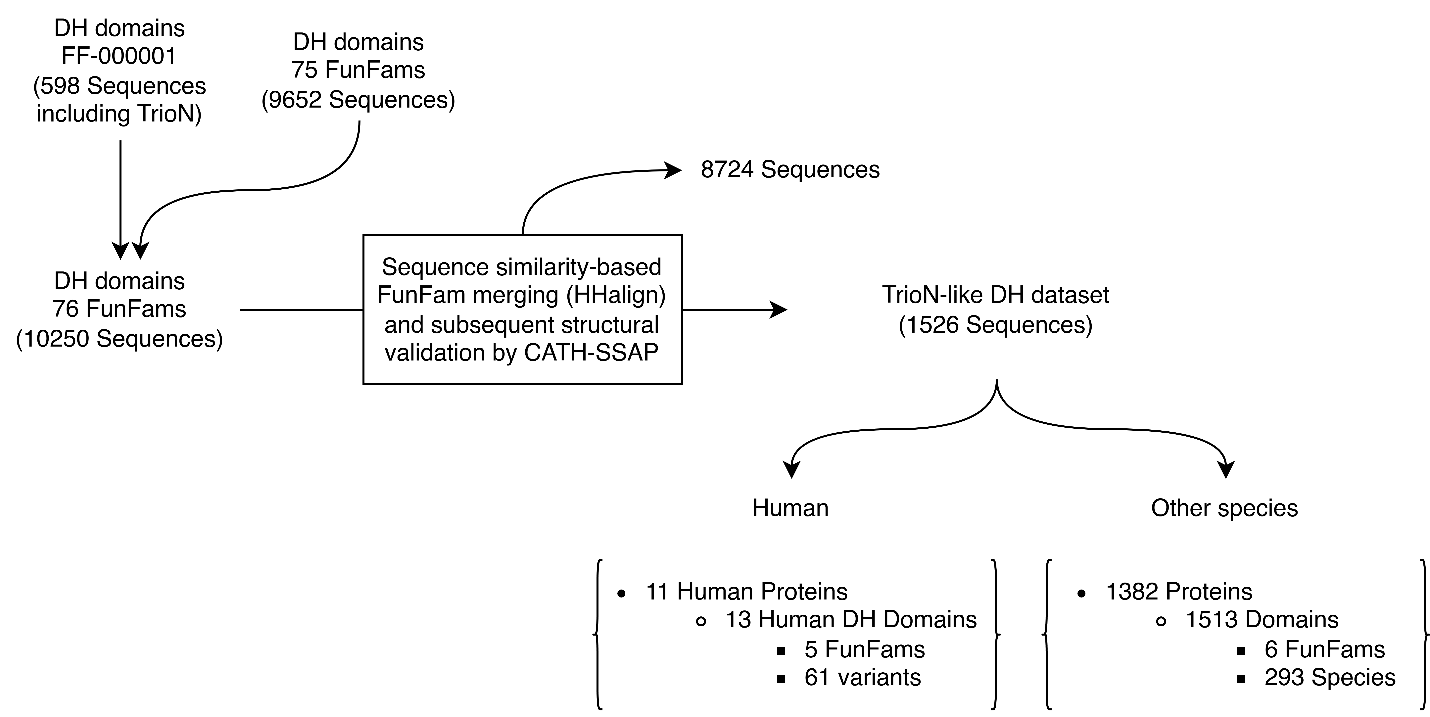

**Supplementary Figure S7:** Process diagram showing the sequence-based and structure-based analyses and the number of sequences and structures at each step. The N-terminal TRIO DH domain (TrioN) is assigned to functional family FF-000001 by the CATH-Gene3D resource. The sequences in FF-000001 were expanded with DH domains from other functional families that have a high probability of being structurally and functionally similar to TrioN using a selection protocol consisting of sequence-based analysis (HHalign) and structural analysis (CATH-SSAP). See text.

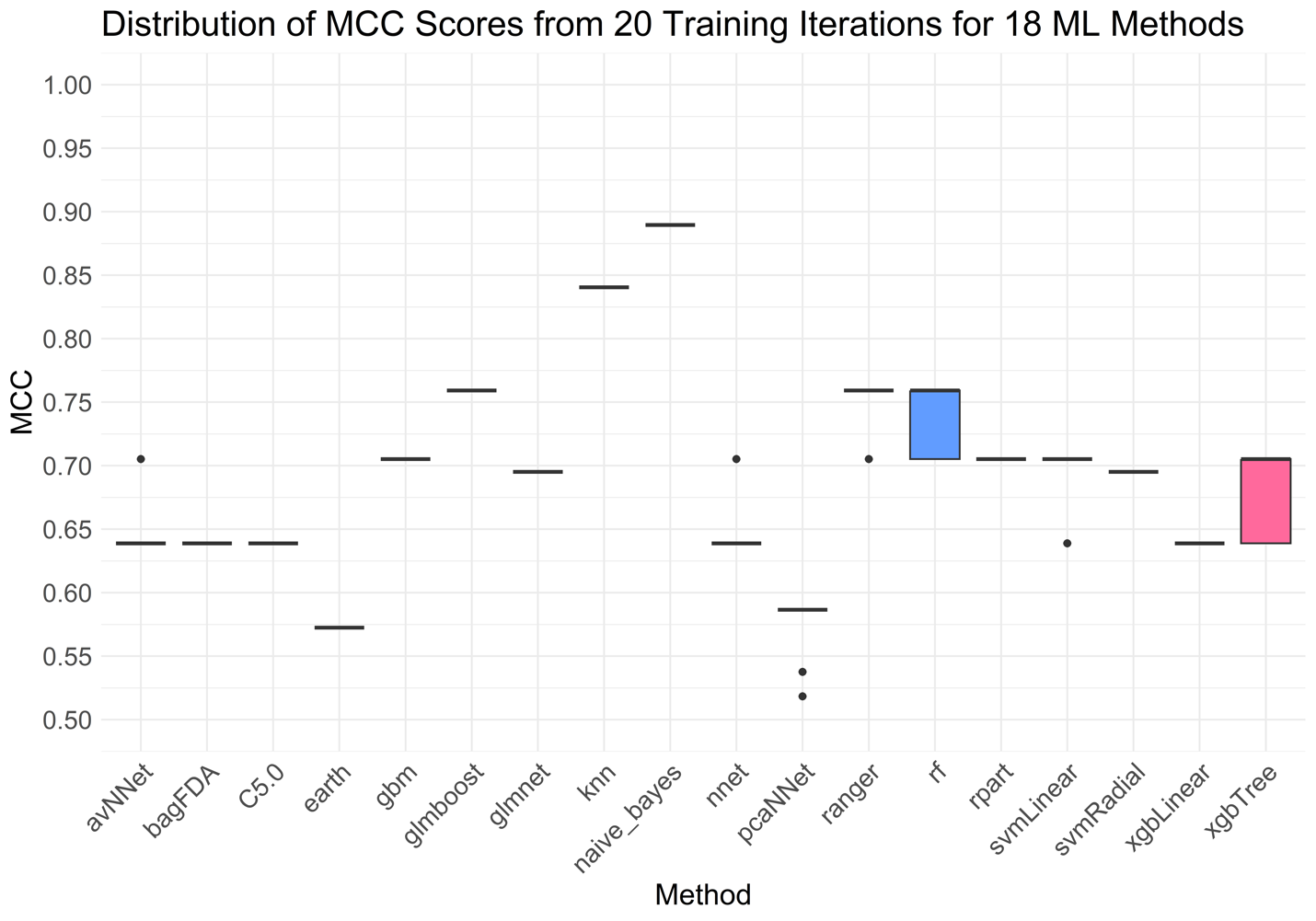

**Supplementary Figure S8:** Performance of 18 different machine learning methods was measured using the MCC metric on the TrioN DH dataset. Each method was trained 20 times. All features (Formula: Inclusive) in TrioNsight methodology are included in the training. avNNet, nnet, pcaNNet, ranger, random forest, svmLinear and xgbTree methods had varying performances across training iterations, while all other methods had fixed performances.

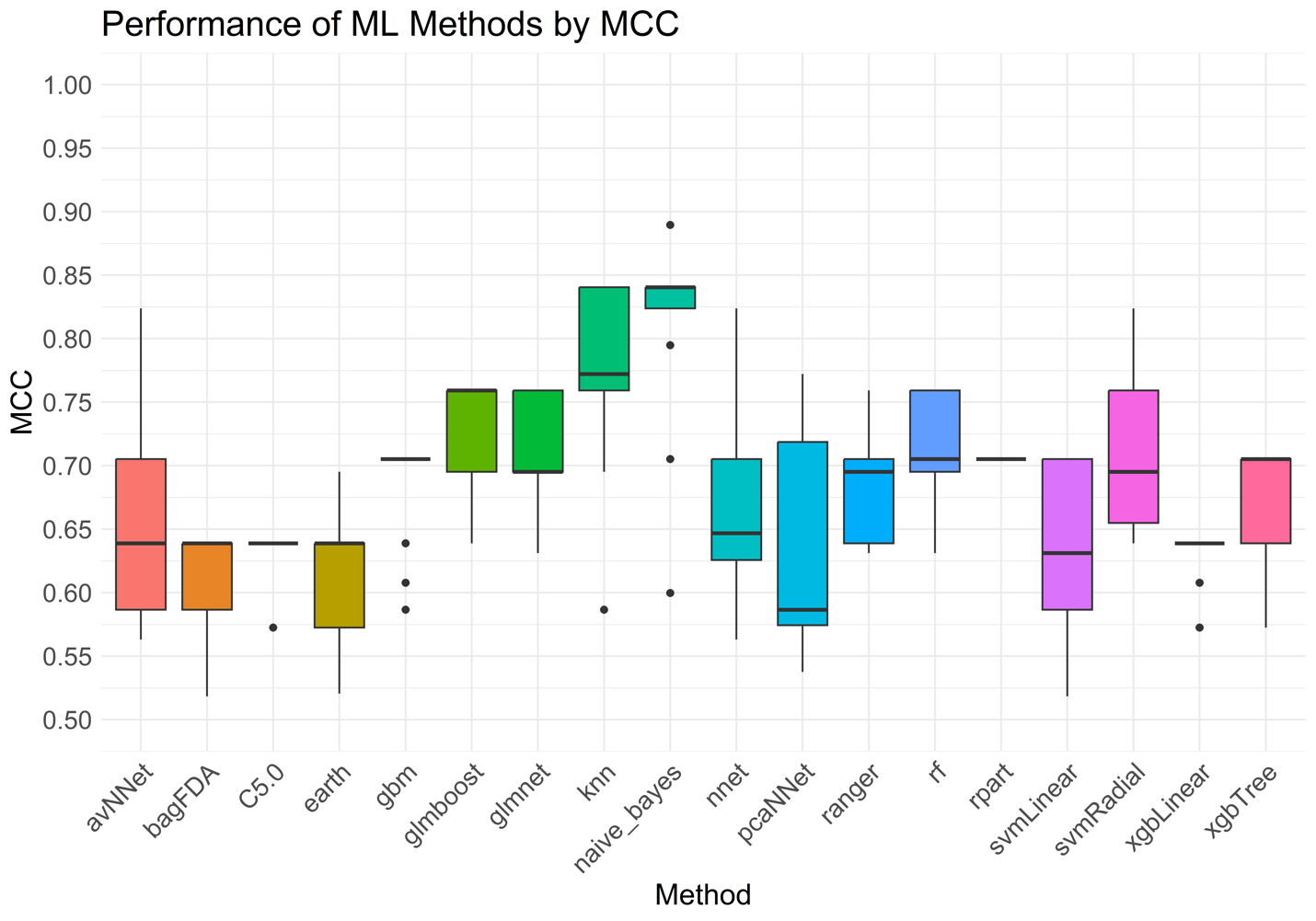

**Supplementary Figure S9:** 18 different machine learning methods were trained 13 times using different feature sets (see Figure 9) on the TrioN DH dataset and performances were measured using the MCC metric. Apart from the formula ‘Inclusive’, which contains all features in the TrioNsight methodology (see Figure 4), other formulas leave one or more (i.e., the formula ‘w/o External Predictors’) features out of the training. The aim is to determine the most robust machine learning method for the change in features. Based on this result, naïve-Bayes is the most robust method with a higher MCC score and low variance compared to other methods.

**Supplementary Table S1:** Complete list of options evaluated to find the best method with which to train TrioNsight. The caret R library implementations of these methods were applied to the TrioN-like DH dataset and performances were measured. The best-performing method was selected to continue with the feature selection step.

| 1. Rf | 2. svmRadial | 3. svmLinear | 4. gbm | 5. glmnet |
| --- | --- | --- | --- | --- |
| 6. glmboost | 7. knn | 8. nnet | 9. avNNet | 10. naïve_bayes |
| 11. xgbTree | 12. xgbLinear | 13. ranger | 14. earth | 15. pcaNNet |
| 16. rpart | 17. C5.0 | 18. bagFDA |  |  |

**Supplementary Table S2** (See file Supplementary Table S2.xlsx): Variant impact predictions for TrioN-like DH domain residues. The table presents meta-predictor scores for 50,122 variant predictions across 13 DH domains within the TrioN-like dataset. For each residue position, 19 potential amino acid substitutions were evaluated, representing the probable missense variants. Predictions include both previously reported variants from existing databases and novel hypothetical variants.

**Supplementary Table S3** (See file Supplementary Table S3.xlsx): Table presenting the complete dataset of TrioN-like DH domains used in this study, including domain sequences, structural boundaries, functional classifications, and source protein information with taxonomic annotations.

**Supplementary Table S4** (see file Table Supplementary S4.xlsx): Table presenting structural similarity of DH domains across all the paralogs based on distribution of RMSD values and SSAP scores. All models were aligned to the representative TrioN model using the CATH-SSAP algorithm. The representative TrioN model is shown in the first row of the table. The table contains proteins, domain boundaries, domain lengths, aligned residues from the target domain, the percent amino acid identity based on the alignment, SSAP scores, and equivalenced percentages which are based on the ratio of aligned residue to the total target domain length, and RMSD scores.

**Supplementary Table S5:** Determining the best clustering results with a ꭓ2 test applied to all strucclus runs n from 2 to 8 (on the left). The best p-value is shown in bold. On the right, variant classes in each cluster for n=5 along with the size of each cluster.

| Number of Clusters | p-value | ꭓ^2^ |  | Clusters | Benign | Pathogenic | Cluster Size |
| --- | --- | --- | --- | --- | --- | --- | --- |
| 2 | 0.3661 | 0.8168 |  | Cluster 1 | 16 | 5 | 21 |
| 3 | 0.0013 | 13.2464 |  | Cluster 2 | 1 | 9 | 10 |
| 4 | 0.0024 | 14.3745 |  | Cluster 3 | 9 | 10 | 19 |
| 5 | 0.0020 | 16.9413 |  | Cluster 4 | 1 | 7 | 8 |
| 6 | 0.0039 | 17.3232 |  | Cluster 5 | 2 | 1 | 3 |
| 7 | 0.0070 | 17.7242 |  |  |  |  |  |
| 8 | 0.0081 | 19.0337 |  |  |  |  |  |

**Supplementary Table S6** (see file Table S6.xlsx): Performance comparison of 18 machine learning methods evaluated for TrioNsight training through systematic feature ablation. Methods were assessed under different feature sets: ‘Inclusive’ models contained all available features; ‘w/o [feature]’ models excluded individual features during training; ‘Exclusive’ models omitted all variant impact prediction algorithms (AlphaMissense, VariPred, PHACT, and PHACTboost); ‘CompPredictorsOnly’ models functioned as meta-predictors utilising exclusively the four variant impact prediction algorithms as input parameters.

**Supplementary Table S7:** Performance was assessed using seven metrics to compare TrioNsight against eight established variant impact prediction algorithms across the complete dataset, including overlapping variants. Comparator algorithms included CADD, ESM-1b, AlphaMissense, VariPred, MutPred2, PolyPhen-2, PHACT, and PHACTboost. Best-performing scores for each metric are highlighted in bold.

| Predictor | Precision | Recall | Accuracy | False Positive Rate | False Negative Rate | F1 Score | MCC |
| --- | --- | --- | --- | --- | --- | --- | --- |
| CADD | 0.871 | 0.844 | 0.853 | 0.138 | 0.156 | 0.857 | 0.705 |
| ESM-1b | 0.775 | 0.969 | 0.836 | 0.310 | 0.031 | 0.861 | 0.692 |
| AlphaMissense | 0.879 | 0.879 | 0.869 | 0.143 | 0.121 | 0.879 | 0.736 |
| VariPred | 0.963 | 0.813 | 0.885 | 0.035 | 0.188 | 0.881 | 0.782 |
| MutPred2 | 0.833 | 0.938 | 0.869 | 0.207 | 0.063 | 0.882 | 0.742 |
| PolyPhen-2 | 0.732 | 0.938 | 0.787 | 0.379 | 0.063 | 0.822 | 0.594 |
| PHACT | 0.048 | 0.031 | 0.164 | 0.690 | 0.970 | 0.038 | -0.692 |
| PHACTboost | 0.861 | 0.969 | 0.902 | 0.172 | 0.031 | 0.912 | 0.809 |
| TrioNsight | 1.000 | 0.906 | 0.951 | 0.000 | 0.094 | 0.951 | 0.906 |

**Supplementary Movie S1** (see file Supplementary Movie S1.mp4): Rotational visualisation of amino acid conservation across different DH domain sets mapped onto the representative TrioN structure (1NTY - DH domain coordinates). The animation displays sequence conservation patterns derived from multiple sequence alignments of three distinct sets: A) non-redundant sequences from all DH domain functional families (clustered at 70% sequence identity using MMSeqs2), B) the TrioN functional family (FF-000001), and C) TrioN-like DH domains identified through sequence and structural similarity. The structure is colour-coded according to Scorecons conservation scores, with regions conserved across all three sets highlighted by continuous circular outlines, and regions conserved in only two sets indicated by dashed circular outlines. This dynamic perspective demonstrates how conservation patterns vary across different evolutionary contexts within the DH domain architecture.

**Supplementary Movie S2** (see file Supplementary Movie S2.mp4): Rotational visualisation of variant clusters on the representative 1NTY TrioN-DH model. The animation shows rotational views of five variant clusters identified by the strucclus algorithm on variants inherited across TrioN-like DH domains, with TRIO-Rac1 binding considered in determining the optimal cluster number. Space-filled representations highlight pathogenic variants in red and benign variants in green, demonstrating their spatial distribution and potential functional implications within the protein structure.

### **References**

[1]	Lin W, Wells J, Wang Z, et al. Enhancing missense variant pathogenicity prediction with protein language models using VariPred. Scientific Reports 2024 14:1 2024;14:1–13. https://doi.org/10.1038/s41598-024-51489-7.

[2]	Cheng J, Novati G, Pan J, et al. Accurate proteome-wide missense variant effect prediction with AlphaMissense. Science (1979) 2023;381. https://doi.org/10.1126/science.adg7492.

[3]	Kuru N, Dereli O, Akkoyun E, et al. PHACT: Phylogeny-Aware Computing of Tolerance for Missense Mutations. Mol Biol Evol 2022;39. https://doi.org/10.1093/molbev/msac114.

[4]	Dereli O, Kuru N, Akkoyun E, et al. PHACTboost: A Phylogeny-Aware Pathogenicity Predictor for Missense Mutations via Boosting. Mol Biol Evol 2024;41. https://doi.org/10.1093/MOLBEV/MSAE136.

[5]	Won DG, Kim DW, Woo J, et al. 3Cnet: Pathogenicity prediction of human variants using multitask learning with evolutionary constraints. Bioinformatics 2021;37. https://doi.org/10.1093/bioinformatics/btab529.

[6]	Ioannidis NM, Rothstein JH, Pejaver V, et al. REVEL: An Ensemble Method for Predicting the Pathogenicity of Rare Missense Variants. Am J Hum Genet 2016;99. https://doi.org/10.1016/j.ajhg.2016.08.016.

[7]	Miller DT, Lee K, Abul-Husn NS, et al. ACMG SF v3.1 list for reporting of secondary findings in clinical exome and genome sequencing: A policy statement of the American College of Medical Genetics and Genomics (ACMG). Genetics in Medicine 2022;24. https://doi.org/10.1016/j.gim.2022.04.006.

[8]	Xie X, Moon PJ, Crossley SWM, et al. Oxidative cyclization reagents reveal tryptophan cation–π interactions. Nature 2024;627. https://doi.org/10.1038/s41586-024-07140-6.

[9]	McCallum M, Park YJ, Stewart C, et al. Human coronavirus HKU1 recognition of the TMPRSS2 host receptor. Cell 2024;187:4231-4245.e13. https://doi.org/10.1016/J.CELL.2024.06.006/ASSET/7F9CADC1-15A0-4B9C-9119-079762260E9F/MAIN.ASSETS/GR5.JPG.

[10]	Frazer J, Notin P, Dias M, et al. Disease variant prediction with deep generative models of evolutionary data. Nature 2021;599. https://doi.org/10.1038/s41586-021-04043-8.

[11]	Jagota M, Ye C, Albors C, et al. Cross-protein transfer learning substantially improves disease variant prediction. Genome Biol 2023;24. https://doi.org/10.1186/s13059-023-03024-6.

[12]	Sillitoe I, Cuff AL, Dessailly BH, et al. New functional families (FunFams) in CATH to improve the mapping of conserved functional sites to 3D structures. Nucleic Acids Res 2013;41. https://doi.org/10.1093/nar/gks1211.

[13]	Jumper J, Evans R, Pritzel A, et al. Highly accurate protein structure prediction with AlphaFold. Nature 2021 596:7873 2021;596:583–9. https://doi.org/10.1038/s41586-021-03819-2.

[14]	Lewis TE, Sillitoe I, Dawson N, et al. Gene3D: Extensive prediction of globular domains in proteins. Nucleic Acids Res 2018;46. https://doi.org/10.1093/nar/gkx1069.

[15]	Consortium TU, Bateman A, Martin M-J, et al. UniProt: the Universal Protein Knowledgebase in 2025. Nucleic Acids Res 2025;53:D609–17. https://doi.org/10.1093/NAR/GKAE1010.

[16]	Lees J, Yeats C, Perkins J, et al. Gene3D: A domain-based resource for comparative genomics, functional annotation and protein network analysis. Nucleic Acids Res 2012;40. https://doi.org/10.1093/nar/gkr1181.

[17]	Steinegger M, Meier M, Mirdita M, et al. HH-suite3 for fast remote homology detection and deep protein annotation. BMC Bioinformatics 2019;20. https://doi.org/10.1186/s12859-019-3019-7.

[18]	Das S, Lee D, Sillitoe I, et al. Functional classification of CATH superfamilies: a domain-based approach for protein function annotation. Bioinformatics 2015;31:3460. https://doi.org/10.1093/BIOINFORMATICS/BTV398.

[19]	Friedberg I. Automated protein function prediction - The genomic challenge. Brief Bioinform 2006;7. https://doi.org/10.1093/bib/bbl004.

[20]	Orengo CA, Taylor WR. SSAP: sequential structure alignment program for protein structure comparison. Methods Enzymol 1996;266. https://doi.org/10.1016/s0076-6879(96)66038-8.

[21]	Edgar RC. MUSCLE: multiple sequence alignment with high accuracy and high throughput. Nucleic Acids Res 2004;32:1792–7. https://doi.org/10.1093/NAR/GKH340.

[22]	Valdar WSJ. Scoring residue conservation. Proteins 2002;48:227–41. https://doi.org/10.1002/PROT.10146.

[23]	Liu X, Wang H, Eberstadt M, et al. NMR structure and mutagenesis of the N-terminal Dbl homology domain of the nucleotide exchange factor Trio. Cell 1998;95. https://doi.org/10.1016/S0092-8674(00)81757-2.

[24]	Chhatriwala MK, Betts L, Worthylake DK, et al. The DH and PH Domains of Trio Coordinately Engage Rho GTPases for their Efficient Activation. J Mol Biol 2007;368. https://doi.org/10.1016/j.jmb.2007.02.060.

[25]	DeLano WL. The PyMOL Molecular Graphics System, Version 2.3. Schrödinger LLC 2020.

[26]	Walsh R, Thomson KL, Ware JS, et al. Reassessment of Mendelian gene pathogenicity using 7,855 cardiomyopathy cases and 60,706 reference samples. Genetics in Medicine 2017;19. https://doi.org/10.1038/gim.2016.90.

[27]	Boga I, Sag SO, Duman N, et al. A Multicenter Study of Genotype Variation/Demographic Patterns in 2475 Individuals Including 1444 Cases With Breast Cancer in Turkey. Eur J Breast Health 2023;19. https://doi.org/10.4274/ejbh.galenos.2023.2023-2-5.

[28]	Čalyševa J, Vihinen M. PON-SC - program for identifying steric clashes caused by amino acid substitutions. BMC Bioinformatics 2017;18. https://doi.org/10.1186/s12859-017-1947-7.

[29]	David A, Sternberg MJE. Protein structure-based evaluation of missense variants: Resources, challenges and future directions. Curr Opin Struct Biol 2023;80. https://doi.org/10.1016/j.sbi.2023.102600.

[30]	Caswell RC, Gunning AC, Owens MM, et al. Assessing the clinical utility of protein structural analysis in genomic variant classification: experiences from a diagnostic laboratory. Genome Med 2022;14. https://doi.org/10.1186/s13073-022-01082-2.

[31]	Stephenson JD, Totoo P, Burke DF, et al. ProtVar: mapping and contextualizing human missense variation. Nucleic Acids Res 2024;52:W140–7. https://doi.org/10.1093/NAR/GKAE413.

[32]	Grimm DG, Azencott CA, Aicheler F, et al. The evaluation of tools used to predict the impact of missense variants is hindered by two types of circularity. Hum Mutat 2015;36. https://doi.org/10.1002/humu.22768.

[33]	Livesey BJ, Marsh JA. Interpreting protein variant effects with computational predictors and deep mutational scanning. DMM Disease Models and Mechanisms 2022;15. https://doi.org/10.1242/DMM.049510.
