## Supplementary figures and images for "TrioNsight: Building a meta-predictor to evaluate the clinical impact of TrioN-like Dbl-homology domain variants"

### Supplementary Figure S1.png

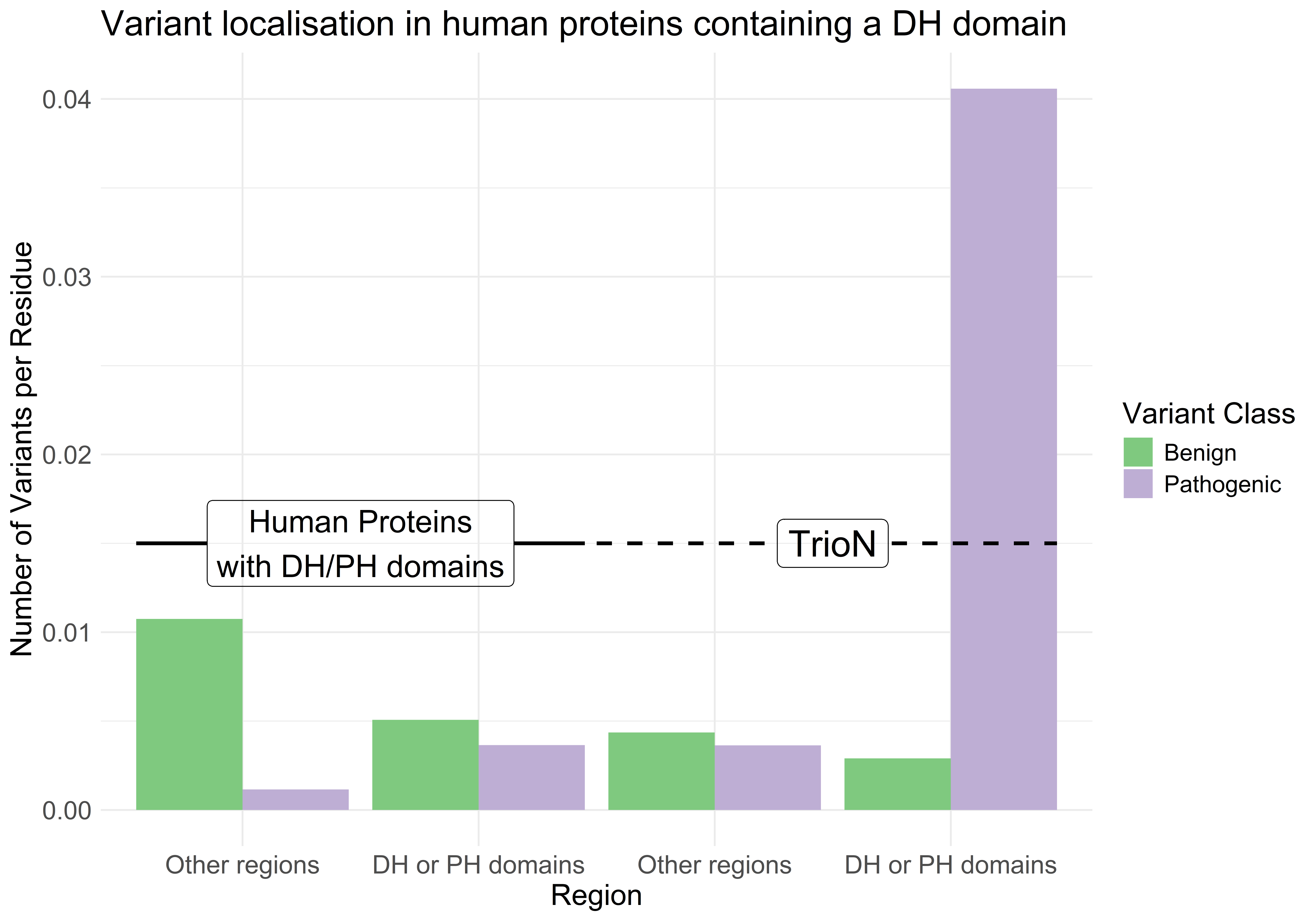

### Supplementary Figure S2.png

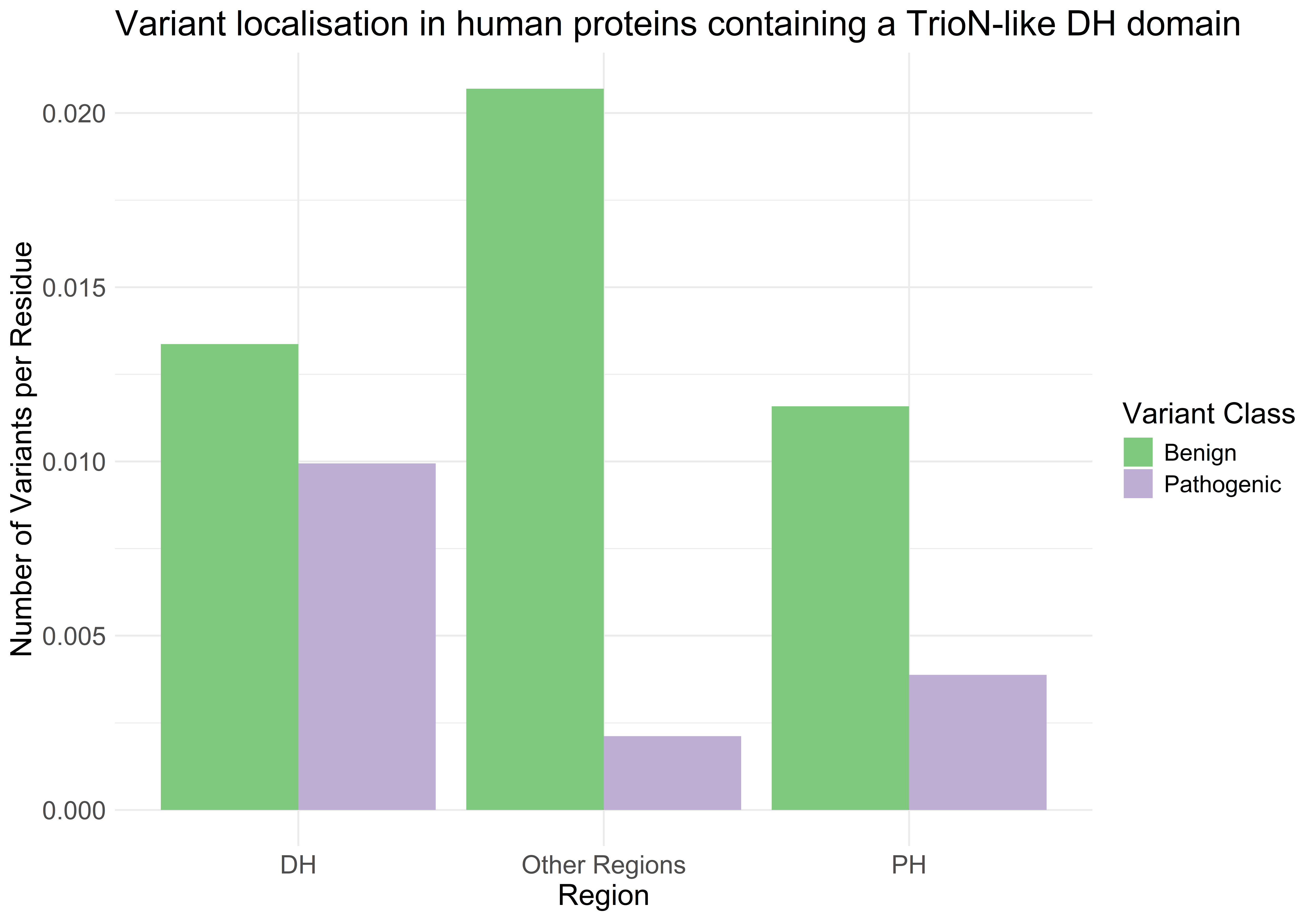

### Supplementary Figure S3.png

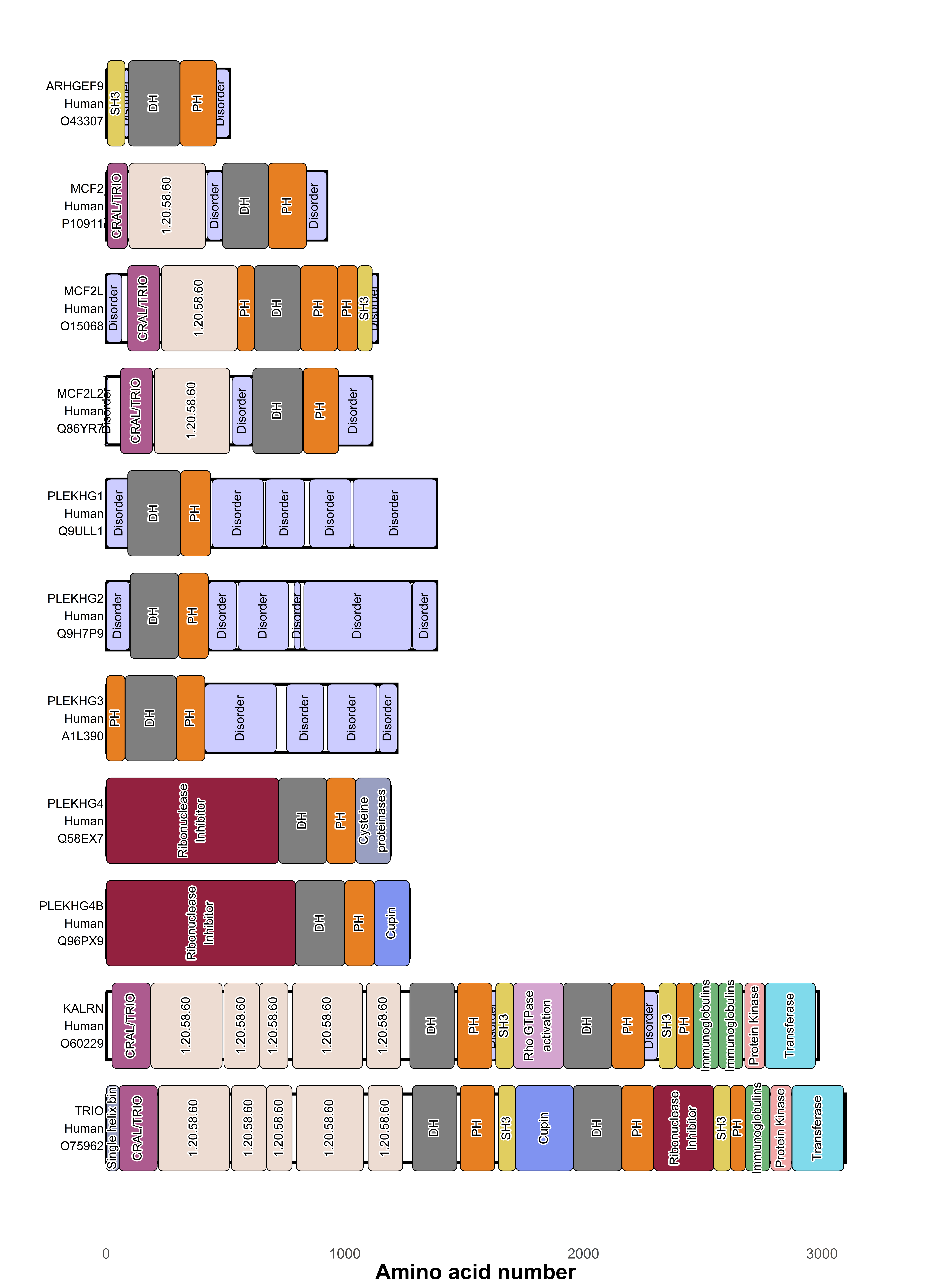

### Supplementary Figure S4.png

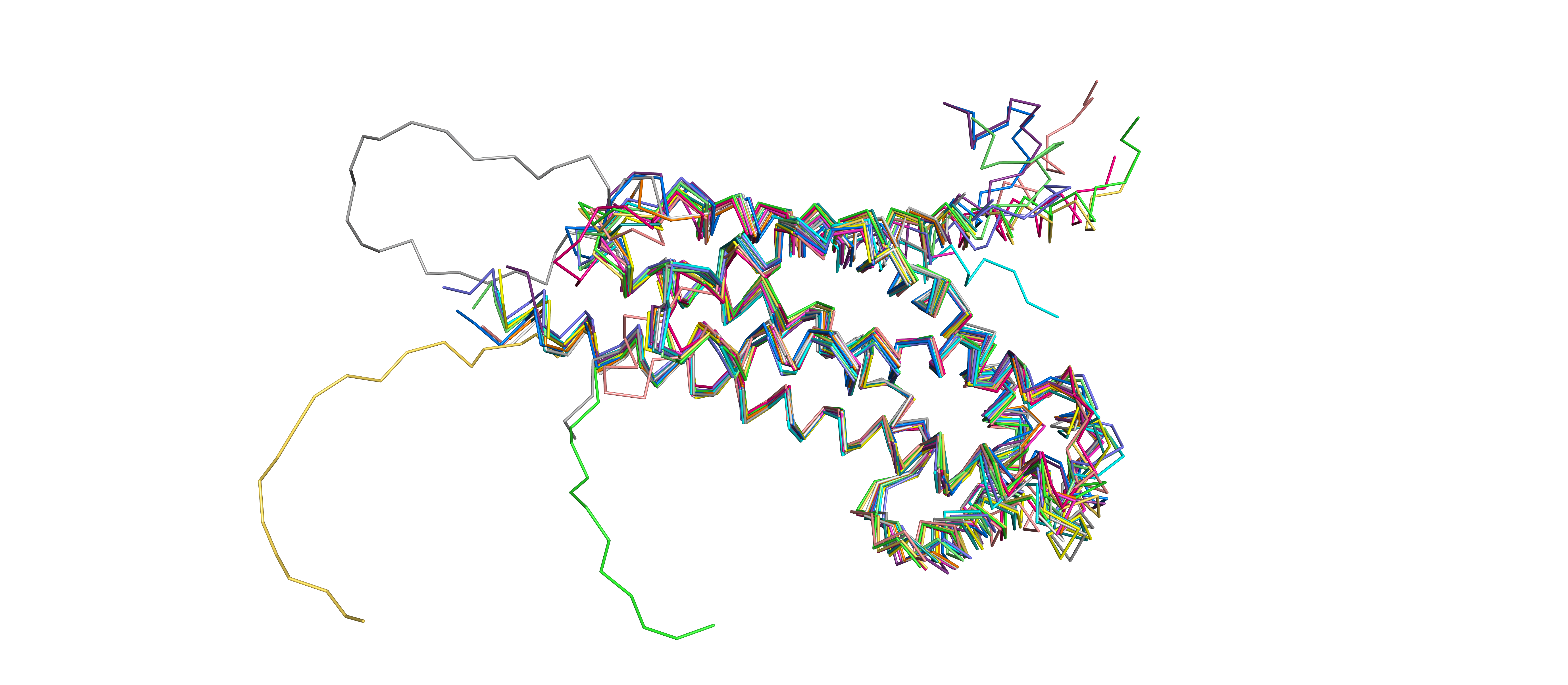

### Supplementary Figure S5.png

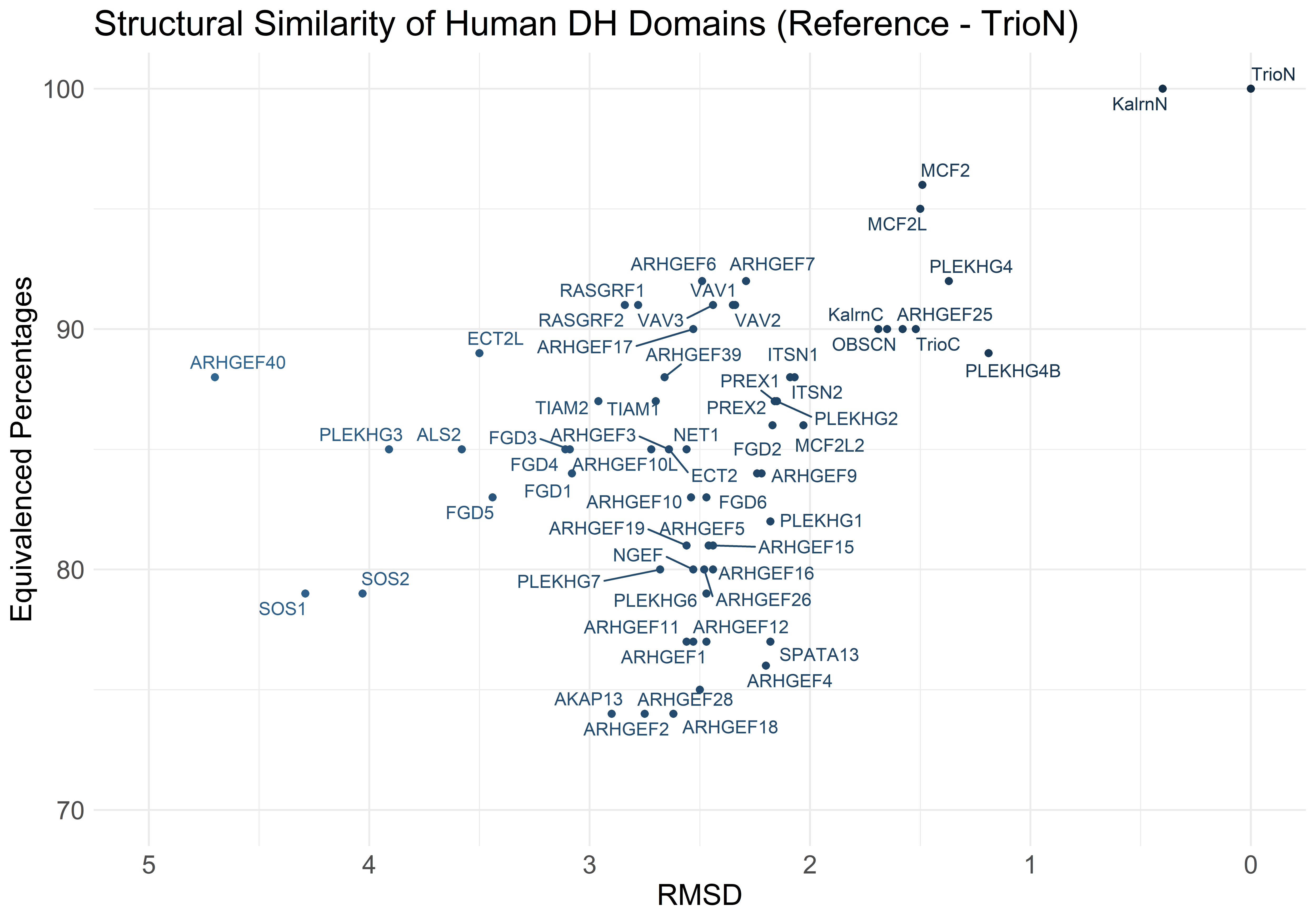

### Supplementary Figure S6.png

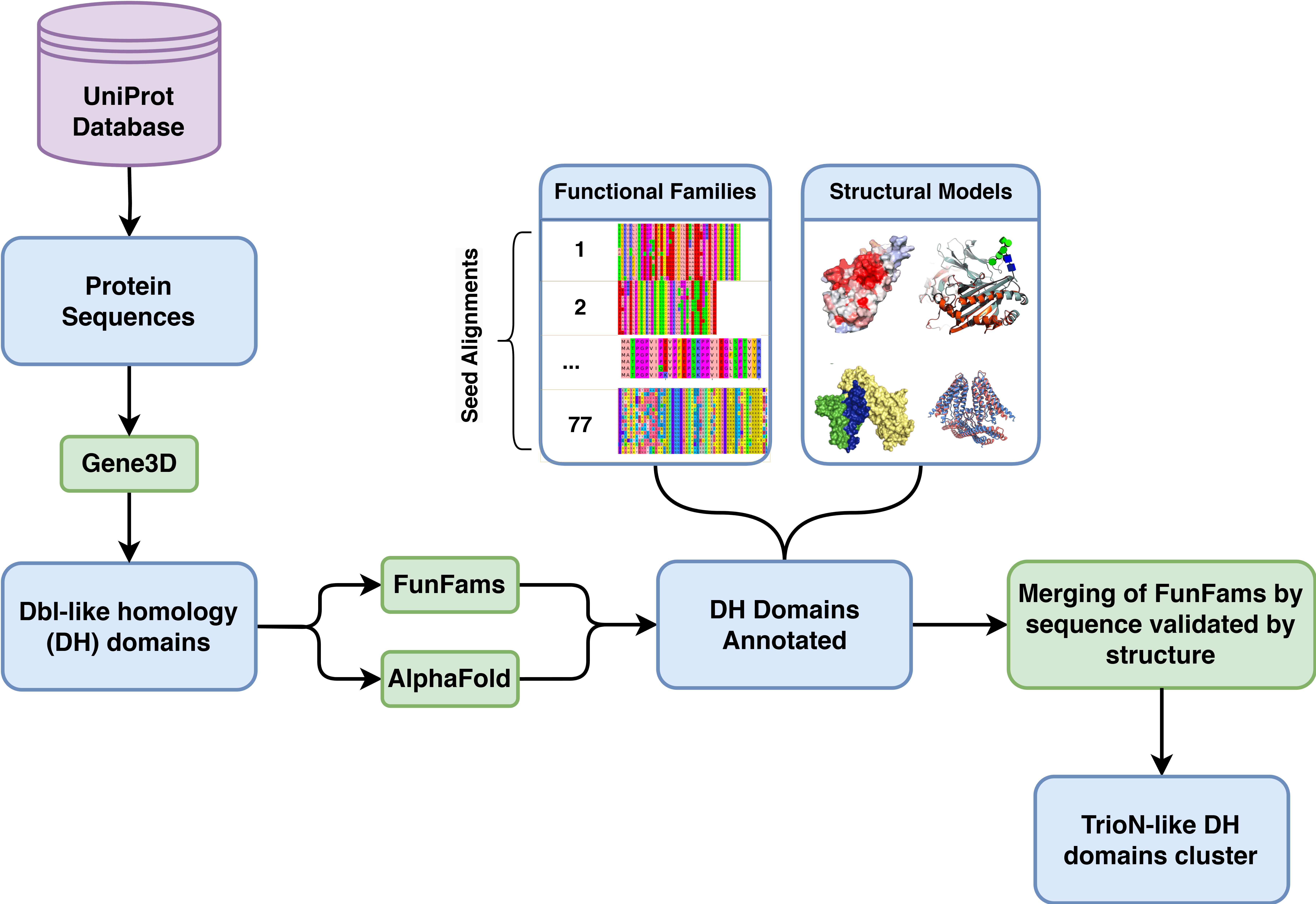

### Supplementary Figure S7.png

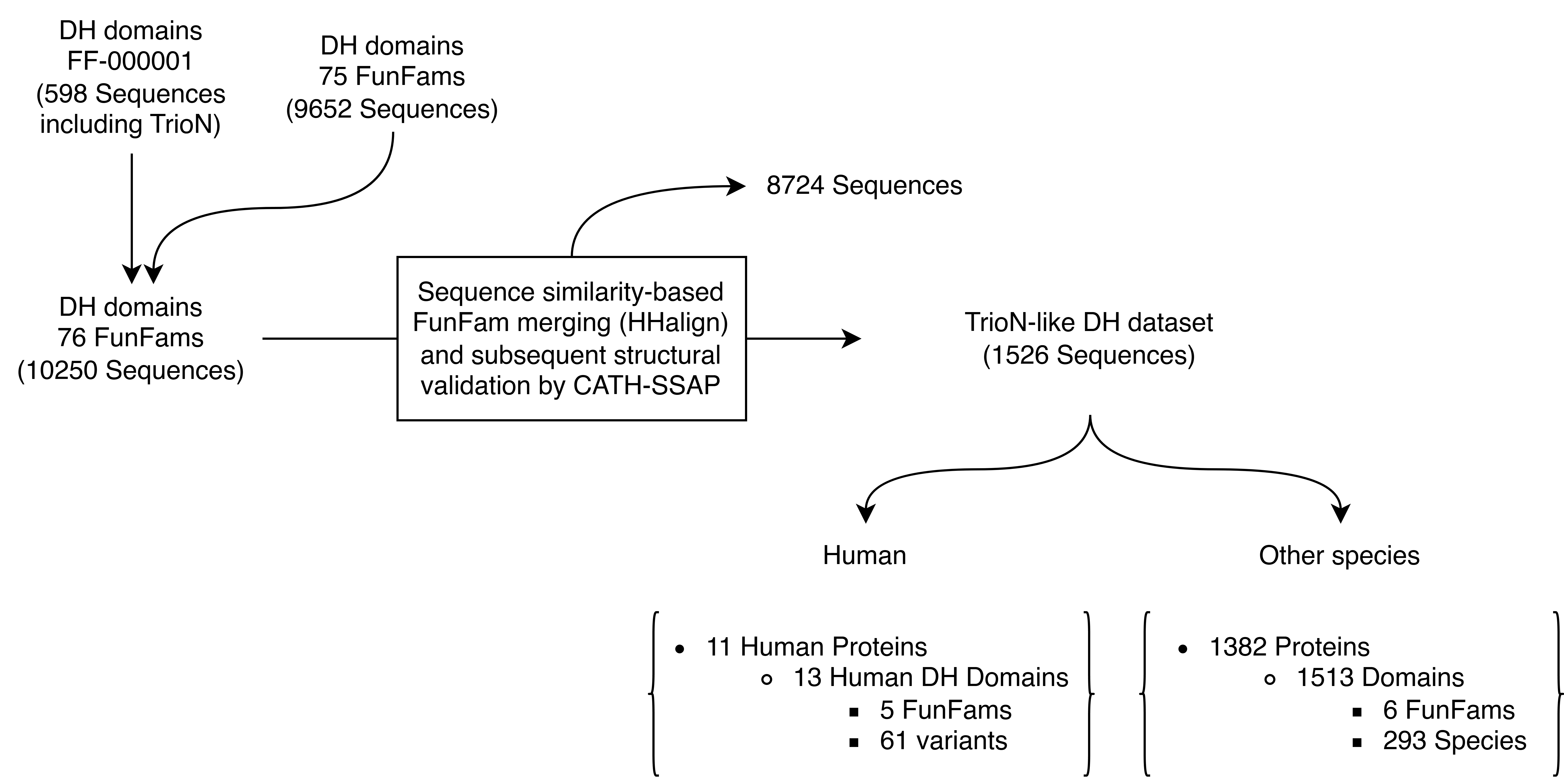

### Supplementary Figure S8.png

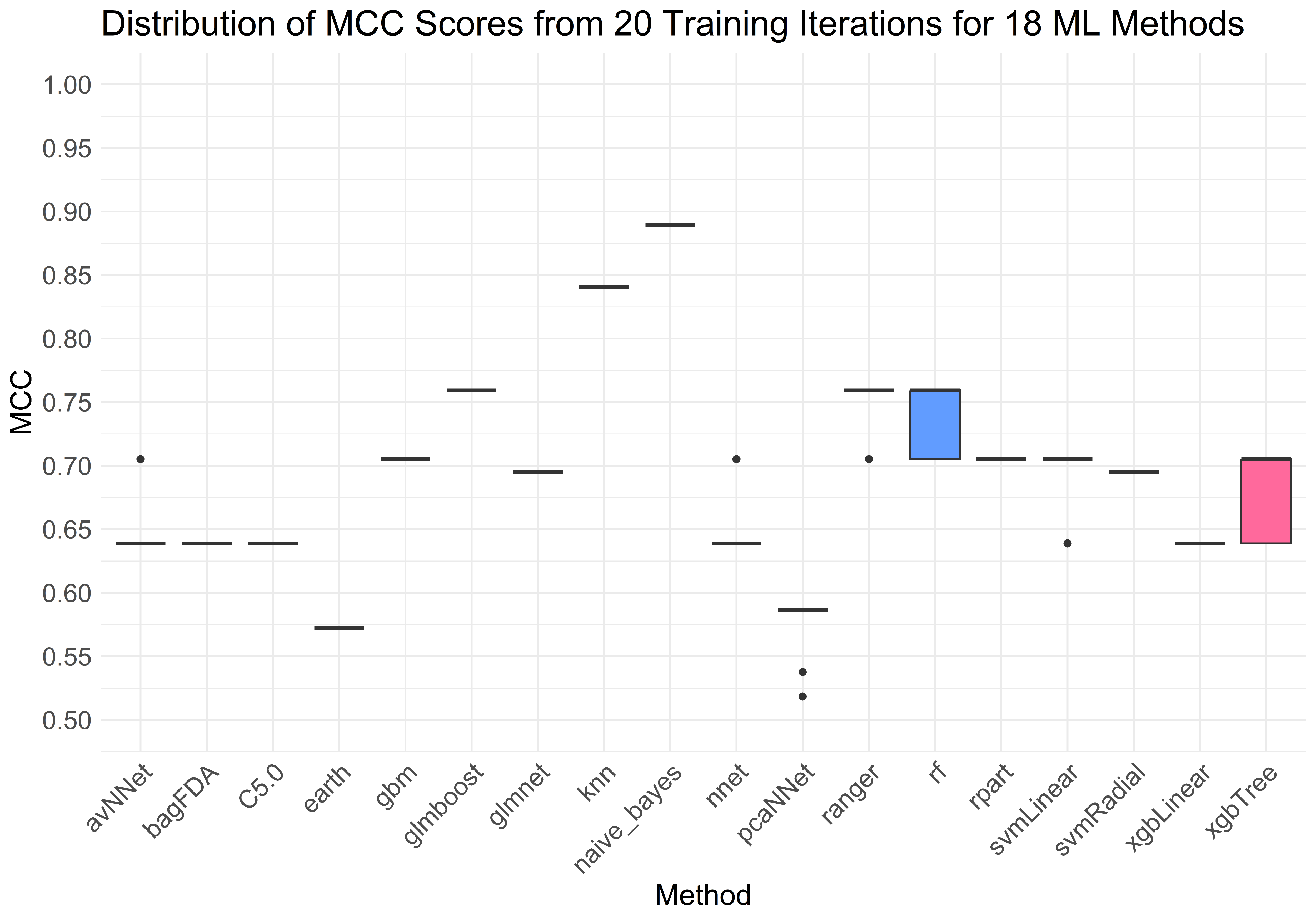

### Supplementary Figure S9.png

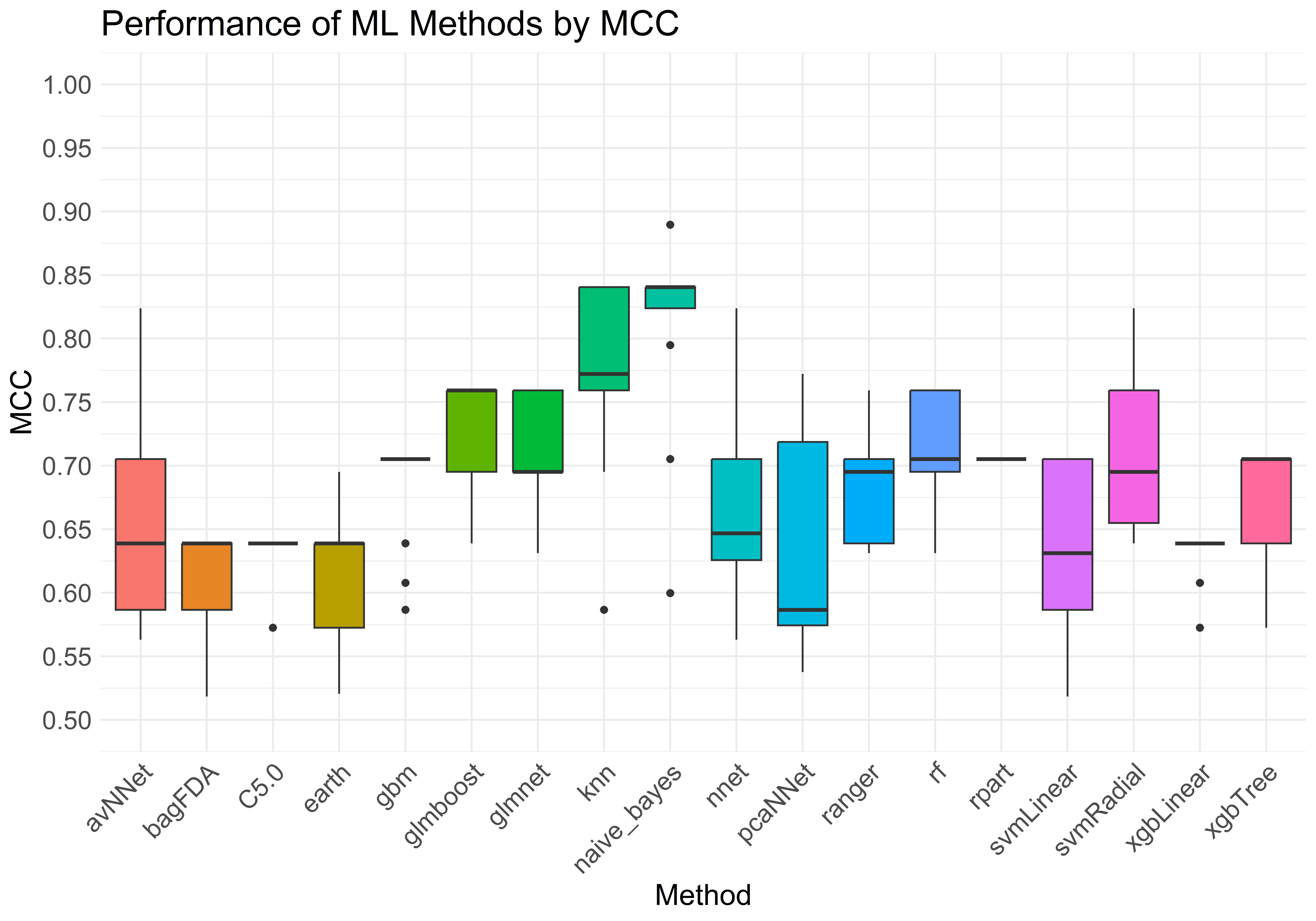
